# Myc and Kras cooperate in adult acinar cells to drive phenotypic heterogeneity, metastasis, and therapeutic resistance in a novel pancreatic cancer mouse model

**DOI:** 10.1101/2025.07.14.664767

**Authors:** Isabel A. English, Patrick J. Worth, Kevin Alexander MacPherson-Hawthorne, Macarena Vergara, Katherine Pelz, Ashley L. Kiemen, Vidhi Shah, Katie E. Blise, Carl Pelz, Motoyuki Tsuda, Michael B. Heskett, Amy T. Farrell, Brittany L. Allen-Petersen, Phillip Jimenez, Meghan Morrison Joly, Mary C. Thoma, Jennifer R. Eng, Colin J. Daniel, Xiaoyan Wang, Melissa Cunningham, Gustavo Salgado-Garza, Jackie L. Phipps, Courtney Betts, Shamilene Sivagnanam, Terry K. Morgan, Laura D. Wood, Lisa M. Coussens, Jonathan Brody, Ellen M. Langer, Rosalie C. Sears

## Abstract

Pancreatic ductal adenocarcinoma (PDAc) is a deadly malignancy, most commonly diagnosed in advanced stages when no curative treatments are available. The development of new models that aid ongoing investigation into the mechanisms by which it initiates, disseminates, and evades treatment is of the utmost importance. *In vivo* models that accurately recapitulate the features and spectrum of human pancreatic cancer are paramount to make a dent in this disease as two decades of the standard-of-care have failed to substantially improve survival. Here, we take advantage of our finding that post-translational stabiliziation of MYC downstream of the canonical PDAc driver, mutant KRAS, is an early event in PDAc progression to design a novel mouse model of PDAc progression based on deregulated, constituitive expression of *Myc* and mutant *Kras* in adult pancreatic acinar cells. Tumors from this KMC model histologically and molecularly recapitulate heterogeneity seen in human PDAc, with a high rate of metastasis to the liver. Cell lines derived from KMC autochthonous PDAc provide new models for orthotopic primary tumors that reliably metastasize to the liver and lung, providing important new tools to efficiently study the metastatic cascade and aid in the develoment of new therapeutics addressing metastatic disease. Cell lines represent distinct molecular subtypes with corresponding differential drug sensitivity. Toghether, this model provides a new and additional tool in the study of pancreatic cancer and the means by which it so deftly evades our best efforts at treatment.

## INTRODUCTION

Pancreatic ductal adenocarcinoma (PDAc) is the third-leading cause of cancer-related deaths and is an aggressive and resistant malignancy.^1,2^ Curative treatment involves both chemotherapy and surgery, unfortunately the majority of patients present with unresectable or metastatic disease and a life expectancy of less than six months.^2^ Current cytotoxic chemotherapeutic regimens have done little to improve disease outcomes or reliably downstage tumors and trials of immunotherapy, more aggressive surgery, and chemoradiotherapy have yielded mixed results often with only incremental progress over the standard-of-care.^3,4^ Furthermore, the promise of RAS inhibition has been overshadowed by rapid development of resistance to these drugs, underscoring the need for agile preclinical model systems that will allow for effective translation to the clinic.^5,6^

*KRAS*, a GTPase-mediator involved in cell signaling, is mutated in 84-95% of pancreatic cancers, leading to its constitutive activation.^7–9^ Diverse *KRAS* mutations are commonly identified in precursor pancreatic intraepithelial neoplasia (PanIN) lesions and are considered an early driver of PDAc.^10^ However, animal models demonstrate that mutant KRAS alone is largely insufficient to drive PDAc.^11–13^ In developing genetically engineered mouse models (GEMM) for the study of pancreatic cancer, secondary drivers involving the loss of key tumor suppressors have been combined with KRAS including *TP53*, *CDKN2A*, and *SMAD4*, all of which are commonly mutated/inactivated in advanced PDAc.^14–17^

KRAS drives stabilization of the MYC oncoprotein via phosphorylation of the Ser62 residue, increasing the cellular expression and activity of MYC and promoting oncogenesis.^18–21^ While rarely mutated itself, increased MYC activity (via amplification, stabilization, and altered degradation) is common in PDAc, and is associated with worse outcomes and metastatic disease.^22–25^ Perhaps more urgently, *MYC* amplifications have been identified as a key mechanism of resistance to KRAS^G12D^ inhibition.^6^

MYC, a key player in numerous hallmarks of cancer, is a major transcriptional regulatory factor that drives cell proliferation, metastasis, and therapeutic resistance.^26^ MYC transcriptional deregulation has been observed early in pancreatic tumorigenesis and is identifiable in precancerous lesions.^27–30^ Similarly, Maddipati *et al.* reported MYC as a major driver in pancreatic liver metastasis. They linked PDAc liver metastasis with MYC signaling activation and showed metastatic patients were enriched for *MYC* gene amplification.^25^ In this study we more thoroughly analyzed the relationship between Ser62 phosphorylation and PDAc development, providing rationale for KRAS-mediated activation of MYC being an early event in tumorigenesis and supporting mouse models of PDAc combining mutant KRAS and MYC.

We previously reported on a GEMM developed using a Pdx1-*Cre* to drive mutant KRAS and deregulated MYC in progenitor cells of the pancreas starting around E7.^22^ In this model, we observed acinar-to-ductal metaplasia (ADM) to PanINs, and progression to invasive PDAc, however we also found high rates of lineage plasticity with mixed adenocarcinoma and neuroendocrine neoplasia.^22^ PDAc is a cancer of older adults with the cell type of origin thought to commonly be the acinar cells.^31–33^ Thus, to better mimic the origins of the human disease we have generated a new GEM model with expression of mutant *Kras^G12D^*and deregulated *Myc* from the *Rosa26* promoter under the control of Ptf1a-*Cre^ERTM^* for inducible recombination and expression specifically in mature acinar cells. This model has yielded several important advances in modeling human PDAc.

We show here that *Kras^G12D^* and deregulated *Myc* expression induced in acinar cells recapitulates human disease progression through early and late precursor lesions with high fidelity. Derived tumors also recapitulated the desmoplasia and histologic features of human PDAc, including immune phenotypes. As in human PDAc, these lesions demonstrate heterogeneity in their transcriptional profiles including both major molecular subtypes and respond variably to treatment; phenotypes that are maintained in cell lines derived from these tumors. Importantly, derived cell lines are reliably metastatic to the liver, one of the most common and deadly metastatic sites, allowing for robust investigation into metastatic disease. Herein we present a novel model for the study of pancreatic cancer with clinically relevant features for the study of the deadliest aspects of this disease.

## RESULTS

### MYC activation is associated with decreased survival and progressive disease in PDAc

*MYC* expression and copy number gain is associated with aggressive tumor behavior and metastasis in PDAc.^7,25,34^ Our lab has produced extensive work on how non-genomic, post-translational modifications—such as phosphorylation at the serine-62 (pS62) residue—influence MYC protein expression and activity.^19–21,35–38^ We have demonstrated that kinases downstream of KRAS directly phosphorylate S62 leading to the accumulation of pS62-Myc that has as increased protein stability and transcriptional activation of target genes. We have generated a serine 62 phosphorylation-specific rat monoclonal antibody to detect cellular MYC activity in both human and mouse formalin fixed paraffin embedded (FFPE) tissue.^39,40^

To explore relative levels of activated MYC across disease stages, we performed immunofluorescence (IF) on FFPE tissue sections of *n* = 8 PDAc patients’ samples with matched adjacent normal and precursor tissues (Figure 1A). We measured increased pS62-MYC intensity across disease stages with progression from normal, to areas of acinar-to-ductal metaplasia (ADM), pancreatic intraepithelial neoplasms (PanINs), and frank PDAc. Significant differences in pS62-MYC intensity between adjacent normal tissue and early grade PanINs (*p =* 0.001) and high-grade PanINs (*p* = 0.047) were identified, supporting a role of MYC in early-stage disease progression (Figure 1B)

**Figure 1:**
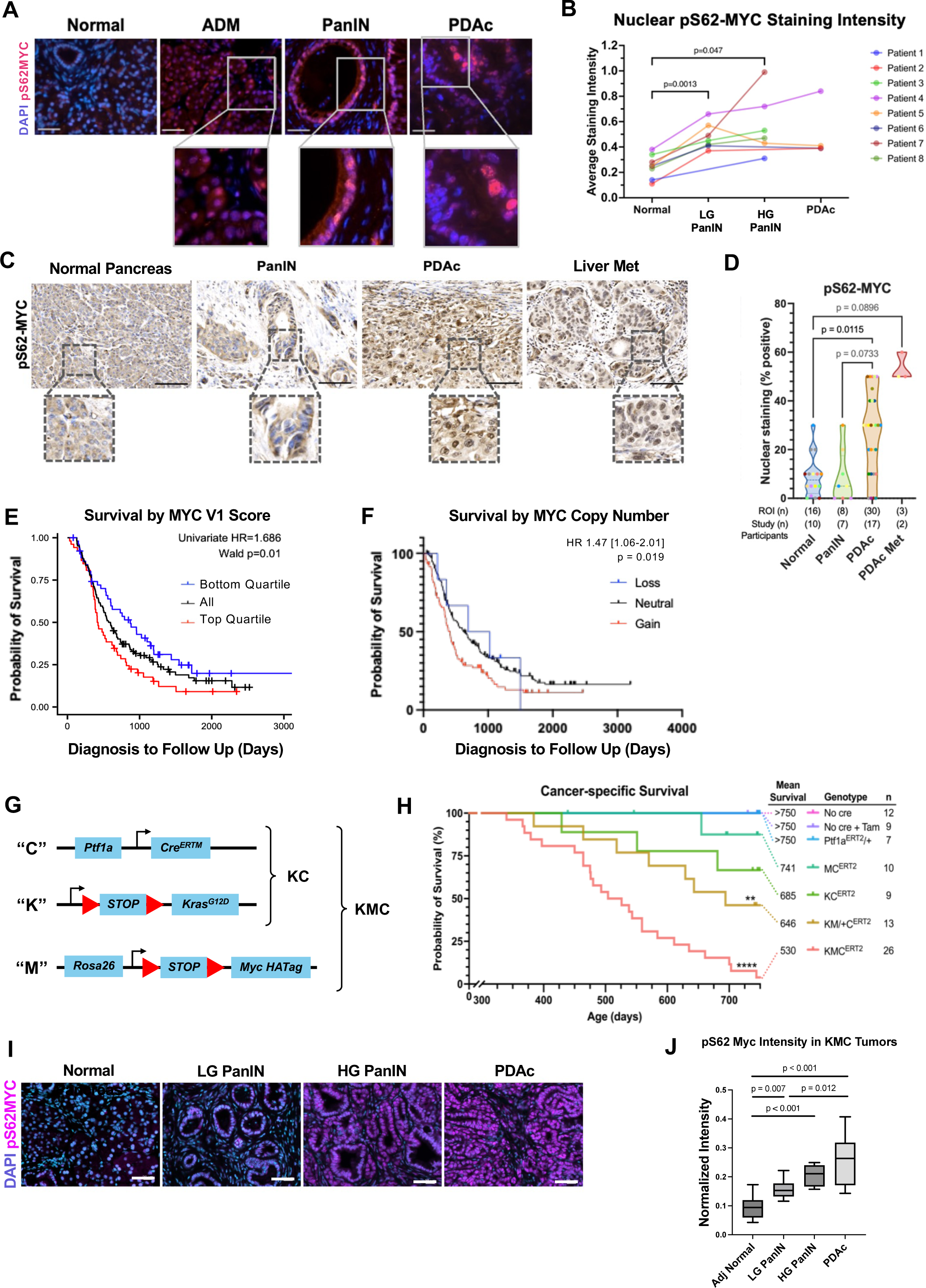
MYC activity increases across progression of human pancreatic cancer and with mutant KRAS is associated with worse outcomes in a novel mouse model. **A.** Representative merged IF of DAPI (blue) and pS62 MYC (pink) on human pancreata demonstrating increased intensity of staining from adjacent normal, ADM, PanIN, and PDAc with magnified insets (scale bars 50µm) **B.** Graph demonstrating quantification of average pS62-MYC nuclear staining intensity by histology (10 randomly identified cells over 5-10 ROIs); evaluated by paired t-tests with Benjamini-Hochberg FDR correction. **C.** pS62-MYC IHC representative images from the “Progression” TMA evaluating nuclear positivity of tumor epithelial cells as scored by percent positive over background. **D.** Graph comparing levels of nuclear staining assessed across disease progression; evaluated by two-way ANOVA with Dunnett’s multiple comparison correction. **E.** Kaplan-Meier curve illustrating overall survival of all patients (black line) compared to the top (red) and bottom (blue) quartile of Hallmarks MYC V1 activity score; n = 208 pts. Cox proportional hazards model. And **F.** by inferred MYC copy number alteration (≥3) in the same population, by Cox proportional hazards model, accounting for LVI, N+, stage, grade, and KRAS mutation. **G.** Gene level schema for the KMC model **H.** Kaplan-Meier curve of cancer specific survival by genotype for the KMC model. Censored animals indicated by tick mark. Significance in median survival compared to KC genotype indicated by asterisk (**p = 0.01; *** p < 0.001) **I.** Representative images of pS62-Myc (pink) and DAPI (blue) merged IF from endpoint KMC tissue demonstrating in adjacent Normal tissues minimal fluorescence, progressing by stage to highest intensity PDAc and metastases. Scale bars indicate 50µm. **J.** Graph demonstrating quantification of average nuclear pS62-Myc staining intensity by histology (10 randomly identified cells over 5-10 ROIs); evaluated by t-tests

We then generated a tissue micro-array (TMA) of twenty-five patients with resected pancreatic cancer and their adjacent dysplastic and normal tissues. Representative 1.5mm diameter punch sections (two per block) included *n =* 20 adjacent normal tissues, *n =* 7 PanIN lesions (low and high grade), *n =* 17 ductal adenocarcinomas, and *n =* 2 disseminated sites (lymph node, liver metastasis). IHC for pS62-MYC was performed on the TMA and quantified as “positive” on a per-cell basis as percentage with nuclear staining above background (Figure 1C). We observed increased proportion of positive cells with pS62-MYC across disease stages, from normal to metastatic tissue (Figure 1D). Together these results demonstrate that MYC stabilization by Ser62 phosphorylation increases early in PDAc initiation and persistently across disease progression. This finding is in agreement with the early detection of activating KRAS mutations in precursor lesions, which leads to accumulation of stabilized pS62-MYC, and its persistence through disease course.

Consistent with the elevation of active MYC across tissues, analysis of our center’s clinically annotated dataset of bulk RNA-sequenced primary tumors (*n* = 230) and metastases (*n* = 70) demonstrated worse overall survival for patients in the highest quartile of MYC V1 GSVA scores (Figure 1E).^24^ From the same dataset, a Cox proportional hazards model for survival across MYC copy number variation (controlling for KRAS mutation, stage, nodal positivity, lymphovascular invasion, and primary tumor grade) yielded significantly worse overall for patients with any MYC copy number gain (CN ≥ 3; Figure 1F).

### Induction of mutant KRAS and dysregulated MYC in adult mouse acini drives ductal metaplasia and PanINs, progressing to adenocarcinoma in a novel genetic mouse model

We previously demonstrated that combining deregulated *Myc* expressed from an knock-in into the *Rosa26* locus--which is not sufficient for oncogenesis on its own--with other mutant oncogenes drives accelerated cancer progression in mice, including PDAc.^22,37,41–43^ Our previously reported noninducible “KMC” model utilized Pdx1-*Cre* to drive expression of *Kras^G12D^* and deregulated *Myc* in the pancreas^22^, similar to the common KPC mouse model of PDAc.^44^ However, *Pdx1*-*Cre* drives Cre expression in the endoderm of the developing pancreas, giving rise to all cell types of the pancreas and this model had a spectrum of pancreatic lesions including adenocarcinomas and a range of neuroendocrine tumors and carcinomas. Since acinar cells are a common cell of origin for PDAc^22,33,45^, we followed this with a *Ptf1a*-*Cre* model to drive expression of *Kras^G12D^*and *Rosa*-*Myc* since the Ptf1a promoter has more restricted expression in pancreatic bud progenitors that give rise to acinar cells (Supplemental Figure 1A).^46–51^ However, this model also gave rise to mixed adenocarcinoma and neuroendocrine tumors and thus we turned to using the tamoxifen-inducible *Ptf1a-Cre^ERTM^*, to fully restrict Cre expression to mature acinar cells.

Adult mice bearing the *LSL-Kras^G12D/+^* (“K”)^37,41,52^, and *Rosa26-LSL^Myc/Myc^*(“M”) alleles were crossed with those expressing the tamoxifen-inducible *Ptf1a-*Cre*^ERTM^* (“C”) to generate the new “KMC” model (Figure 1G; n.b., mice with heterozygous *Rosa26-LSL^Myc/+^* will be referred to as “KM/+C”). Resulting KMC mice are phenotypically normal at birth and resemble their non-recombined littermates. Of note, homozygous *Kras^G12D^* is known to be embryonic lethal.^53^ This genotype is not observed, but the remaining genotypes were seen with expected Mendelian frequency. After induced recombination with intraperitoneal injection of tamoxifen for five days at seven to eight weeks of age, pancreatic acinar cells from the KMC model constitutively expressed *Kras^G12D^* and mildly supra-physiologic levels of *Myc*, similar to a copy number gain of +1 to +2.

Mice were aged in seven genetic cohorts designed to evaluate the combinatorial effect of oncogenic *Kras* mutations and *Myc* deregulation (Figure 1H). We evaluated mice regularly with transabdominal ultrasound and/or abdominal palpation to track the development of tumors and mice were taken down at predefined humane endpoints or at 750 days. Expression of Ptf1a-*Cre^ERTM^*alone caused no pancreatic abnormalities or carcinoma. One MC mouse developed a lymphomatous tumor of the pancreas at 655 days. This agrees with our prior experience and others that *Rosa* driven Myc dysregulation is insufficient to drive tumorigenesis in several organ sites.^22,37,41–43^ Other reports which show Myc alone is sufficient for the development of carcinomas utilize Myc expression at supraphysiologic levels, much higher than exogenous *Rosa-*driven Myc.^22,30^

One-third of KC mice developed pancreatic tumors. When combined with a single copy of exogenous *Myc*, disease latency was significantly reduced in heterozygous KM/+C animals. Ninety-seven percent of KMC homozygotes developed adenocarcinoma, underscoring the cooperativity between mutant KRAS and dysregulated MYC. Importantly, confining recombination and oncogene expression to the acinar compartment via induction in the adult pancreas mitigated the neuroendocrine heterogeneity seen in the embryonic KMC models (Supplemental Fig 1B^22^). Activated pS62-Myc intensity increased across disease state as well (Figure 1I). Low pS62-Myc levels were observed in adjacent normal tissues (Figure 1I, leftmost panel) with significant increases in nuclear intensity measured across disease state (Figure 1J), mirroring characteristics seen in human PDAc (cf. Figure 1A-D). This further supports the role of pS62-Myc in disease progression in this model.

### The KMC model recapitulates human PDAc progression with high histopathologic fidelity

To characterize disease progression in our model, KC, MC, and KMC mice were euthanized 6- and 10-months after tamoxifen induction. A practicing GI pathologist (T.M.) blinded to cohort and genotype examined H&E sections. At both 6- and 10-months, all MC mice (*n =* 4 per genotype at both timepoints) demonstrated benign pancreas tissue with no evidence of dysplasia (Figure 2A). At both 6 and 10 months, ∼50% of mice expressing mutant Kras-alone (KC) harbored low-grade PanIN lesions (*n* = 2/4 6 month and *n =* 2/5, 10 month time point). KMC mice demonstrated low-grade PanIN lesions starting at six months and by the ten-month timepoint 75% of mice showed LG PanINs. Though these cohorts are limited by size, they are consistent with cooperative development of disease seen in the KMC mice. Affected cells progress with high histologic fidelity through known human disease stages including ADM and PanINs, before developing into various grades of ductal adenocarcinoma (Figure 2B).

**Figure 2:**
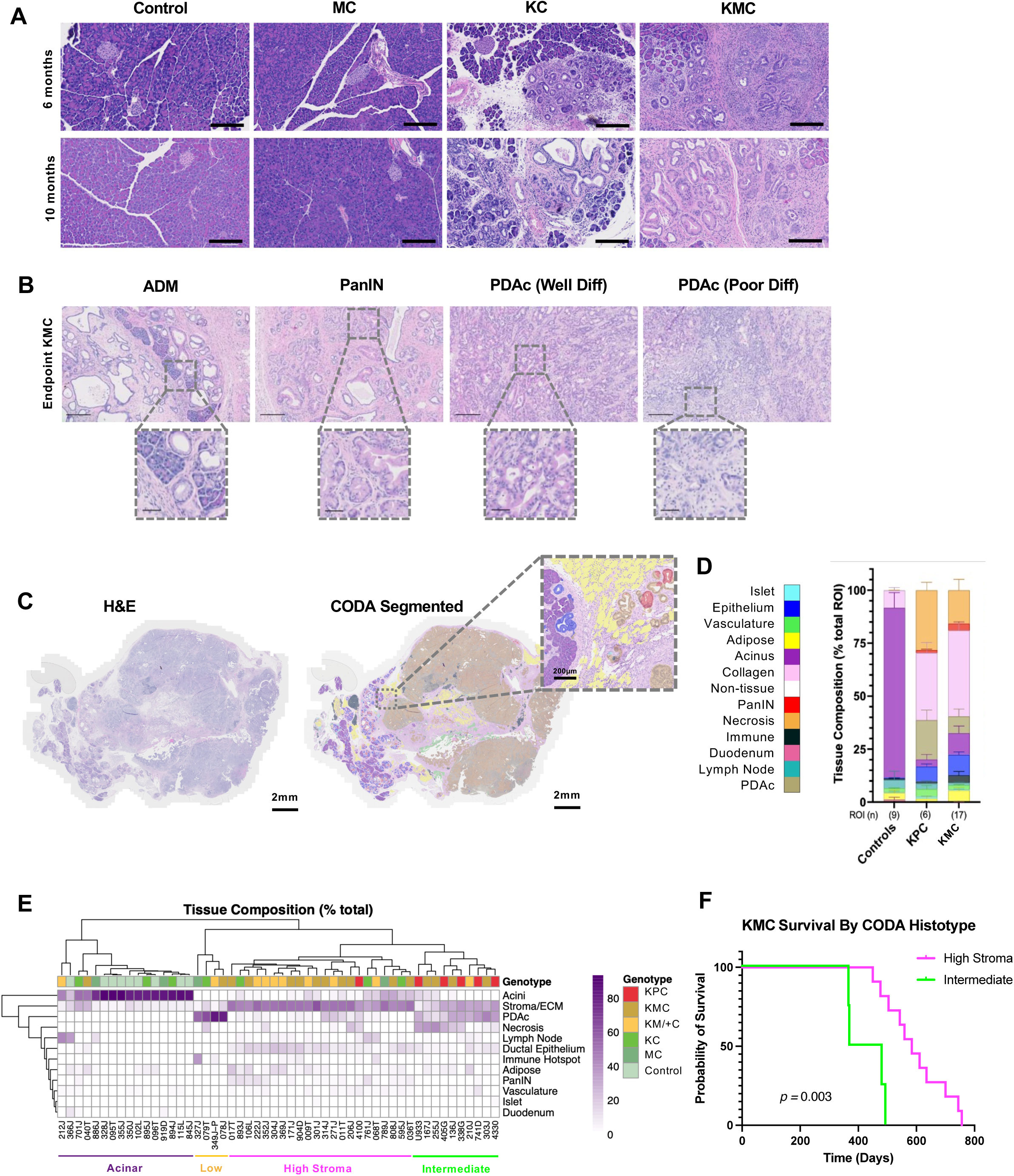
The KMC model recapitulates human PDAc progression with high histopathologic fidelity. **A.** Timepoint tissues by genotype **B.** Histologic progression and types found in endpoint tissues of KMC mice **C.** Representative images of CODA segmentation from whole-section H&E and semantic segmentation. Scale bars indicate 2000µm in whole-slide sections and 200µm in inset. **D.** Summary of weighted percentages of tissue composition for control and KPC vs KMC tumors **E.** Unsupervised clustering based on features identified in each specimen demonstrated clustering reflecting tumor genotype **F.** Kaplan-Meier curve demonstrating survival difference for homozygous KMC mice depending on CODA histologic type

In order to analyze intra- and inter-tumoral histopathologic heterogeneity across the KMC model and genotypes, we utilized CODA, a semantic segmentation deep learning model previously developed for quantification of human pancreas histology by our collaborators at Johns Hopkins University (Fig. 2C)^54^ We analyzed representative H&E sections from nine control, three MC, seven KC, eleven KM/+C, and seventeen KMC endpoint pancreata. Additionally, endpoint tumors from six KPC (*LSL- Kras^G12D/+^*; *LSL*-*Tp53^R172H/+^; Pdx1-Cre*)^44^ mice were included for comparison. The image segmentation algorithm was retrained for mouse histology, and a subset of resulting output was then validated by blinded pathologist (L.D.W.) (supplemental figure 3a) and quantified (Figure 2C inset and 2D, supplemental figure 3b)).

Unsupervised clustering revealed four histologically diverse groups, predominantly separated by their degree of fibrosis (collagen content), a characteristic feature of human PDAc (Figure 2E).^55–58^ We designated these groups as “acinar”, “low-”, “intermediate-”, and “high-desmoplasia”, respectively. ‘Acinar’ histology was primarily composed of KC and MC mice (87.5%) and non-tumorous KM/+C and KMC pancreata (6.3% each). ‘Low-desmoplasia’ tumors had dense tumor cellularity and and included multiple genotypes. High-desmoplasia was the largest group of tumors (*n =* 22, 42.3%) and had marked fibrosis. (Supp. Fig. 6). KMC^”“^ and KM/+C tumors made up the largest percentage of this subtype (50% and 36.3%, respectively). The majority of KPC tumors segregated together as intermediate-stroma, with relatively higher amounts of necrosis seen in this cluster. Stroma-low tumors were diversely composed of cellular PDAc, stroma/ECM, and necrosis, and represented mixed genotypes. In comparison to the evaluated KPC tumors, tissue composition in the KMC model was more desmoplastic and featured more immune aggregates and adipose tissue, though our analysis was underpowered to detect significant differences.

Homozygous KMC tumors clustering in the intermediate desmoplasia group were found to have reduced survival when compared to their high-desmoplasia homozygous KMC counterparts (*p =* 0.003, Figure 2F), indicating biologic as well as phenotypic heterogeneity across this genotype.

### Tumor suppressor loss and genomic instability is a feature of the KMC model

*KRAS* mutation and *MYC* deregulation are early events in PDAc development, but progressive acquisition of other mutations—most commonly in *CDKN2A* (P16), *SMAD4*, and *TP53*—are also a feature of human disease.^14,17,59–62^

To investigate whether the KMC model acquired changes in these genes or their resulting transcripts or proteins, we performed IHC on sections of KMC (*n* = 6) and KPC (*n* = 4) tumors for P53 (Figure 3A) and observed significantly decreased levels of nuclear protein abundance in the KMC model, (Figure 3B). This reduction in protein expression is consistent with the common loss of P53 activity observed in human PDAc and this is associated with poor prognosis in PDAc and other human cancers.^62–66^

**Figure 3:**
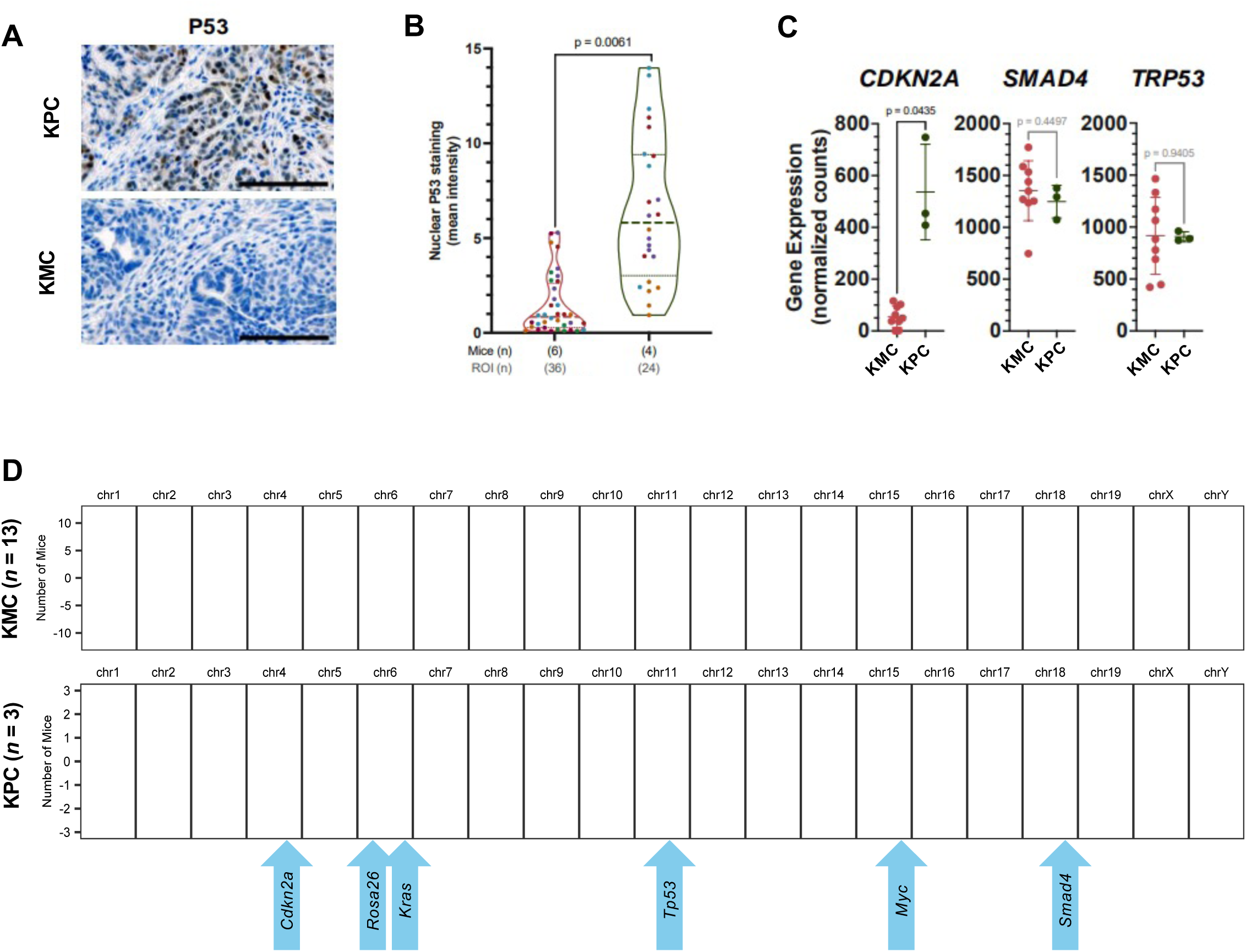
Tumor suppressor loss and genomic instability is a feature of the KMC model. A. Representative IHC demonstrating decreased nuclear staining (brown) in KMC tumors (bottom panel) as compared to KPC tumors (top panel) at endpoint. Scale bars denote 200µm. B. Comparison of percent nuclear staining of KMC vs KPC tumors; each subject identified by dot color. Unpaired t-test with Welch’s correction used to evaluate significance. C. Comparison of normalized gene expression of canonically-mutated genes in PDAc in KMC as compared to KPC tumors; significance calculated using Welch’s ANOVA and Dunnett’s multiple comparisons. D. Genome-level representation of copy number variants by model, annotated with commonly mutated genes in PDAc as well as genes utilized in this model. Blue represents loss; red represents gain.

Endpoint tumors from nine KMC and four KM/+C were subjected to bulk RNA sequencing of tumor enriched regions. Three endpoint KPC tumors were also sequenced as controls and a gold-standard comparator group from tumor enriched regions. We observed a significant decrease in *Cdkn2a* mRNA expression in the KMC tumors when compared to KPC tumors. Both *Smad4* and *p53* transcript levels had high variability in KMC tumors (Figure 3B), but no significant difference in median transcript levels when compared to KPC tumors. Cdkn2a/p16 is known to positively regulate p53 expression in cancers and various tissues.^67^ The observed decrease in *Cdkn2a* transcription may be a mechanism for the decreased nuclear p53 expression observed in the KMC tumors.

We extracted genomic DNA from the same tumor-enriched specimens (above) and performed whole exome sequencing (WES) to probe for spontaneous mutations in high-impact genes. Overall, tumors harbored a low mutational burden (see Supplementary Table 1), and no mutations were identified in canonical PDAc genes *Tp53*, *Cdkn2a* or *Smad4*. This is consistent with recent investigations into other pancreatic cancer GEMMs.^68^

As copy number variation is commonly identified in human PDAc^17,69,70^, we also performed low-pass whole genome sequencing on the above tumors to assess other acquired genomic changes in the models. Alterations in copy number were identified at all three canonical sites (*Tp53*, *Cdkn2a*, and *Smad4*), in addition to *Kras* and *Myc*, demonstrating genomic instability inherent in the KMC model (Figure 3D). The KPC tumors also exhibited copy number alterations; although only 3 tumors were profiled, there appeared less heterogeneity across the tumors.

### Immune heterogeneity and antigen presentation are distinguishing features of the KMC tumor microenvironment

Unsupervised clustering of top and bottom 500 differentially expressed genes in KMC and KPC tumors separated samples by mouse model (Figure 4A). We utilized gene-set enrichment analysis to investigate potential functional differences in expression profiles between the models and found KMC tumors were enriched significantly for nineteen immune-related GO terms (Figure 4B), consistent with the role MYC is known to play in regulating the tumor immune microenvironment.^30,71,72^ Fewer genes were significantly upregulated in KPC tumors and no gene ontology gene sets were significantly enriched in KPC over KMC (Supplementary Figure 5A).

**Figure 4:**
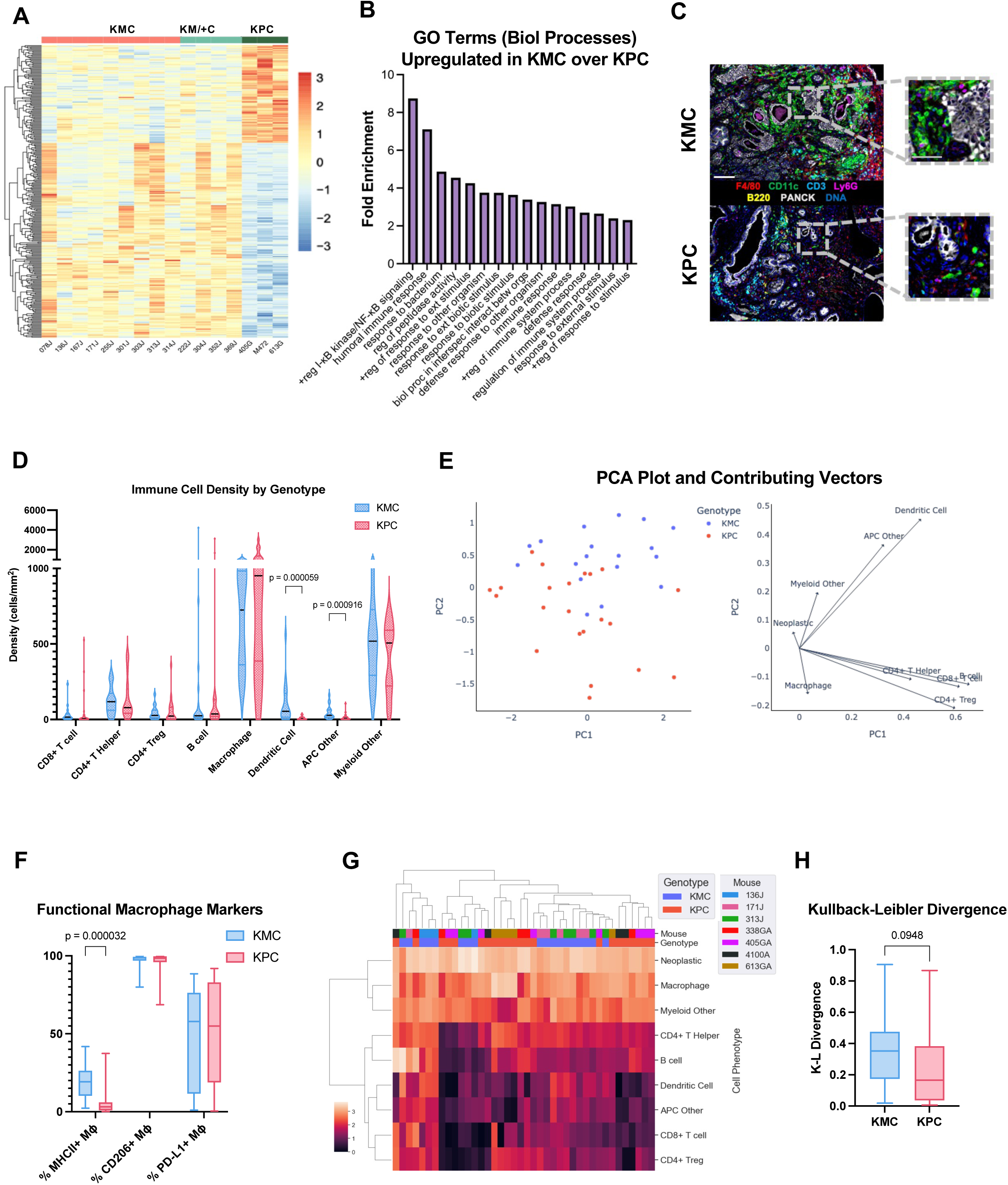
Immune heterogeneity and antigen presentation are distinguishing features of the KMC tumor immune microenvironment. A. Heatmap of KMC, KM/+C, and KPC gene expression clustering B. Gene Ontogeny Terms enriched in KMC transcriptomes versus KPC by differential expression analysis C. Representative image of pseudocolored multiplex IHC of KMC and KPC tumors showing differences in immune microenvironment D. Box plot showing immune cell densities split by genotype. Each dot represents one tissue region (n= 18 KMC, n= 22 KPC). E. Left: PCA showing each tissue region (n=40) plotted according to its log10+1 transformed cell densities and colored according to the genotype of the tumor from which it was sampled from. Right: PCA loadings showing the weights and features contributing to each principal component. F. Box plot showing the percent of macrophages positive for MHCII, CD206, and PD-L1 functional markers split by genotype. Each dot represents one tissue region (n=18 KMC, n= 22 KPC). G. Unsupervised hierarchical clustering of tissue regions (n=40) by the log10+1 transformed cell densities. Top two rows are colored by the mouse from which the tissue region originated and the genotype of the tumor. H. Box plot showing the KL divergence of each tissue region split by genotype (n= 18 KMC, n=22 KPC).

We sought to deeply characterize the KMC microenvironment and immunologic differences suggested by the differential gene expression. To do this, we performed multiplex immunohistochemistry (mIHC) to assess immune contexture within the tumor microenvironments (TMEs) of KMC (*n* = 3) versus KPC (*n* = 4) tumors (see Table 3 for staining sequence and antibodies). Four to eight tissue regions were sampled per tumor, and cells were phenotyped as neoplastic or one of eight immune cell lineages, including lymphoid and myeloid populations (Figure 4C – pseudo-colored images; gating schema in Table 4). Cell densities were then compared between KMC and KPC tumors. While there were no significant differences in total immune cell density or neoplastic cell density between tumor genotype (Supplementary Figure 5B), we did detect significantly higher densities of dendritic cells and APC other cells in KMC tumors than KPC tumors (Figure 4D). Principal component analysis (PCA) further confirmed these results, as indicated by the loadings on the PCA (Figure 4E). Higher densities of dendritic cells, APC other cells, and myeloid other cells were associated with tissue regions from KMC tumors, while higher densities of macrophages and CD4^+^ Tregs were associated with tissue regions from KPC tumors.

To further interrogate the potential function of the immune cells associated with KMC or KPC genotypes, we next assessed expression of specific functional markers on the immune lineages associated with each tumor genotype. While we did not have data to assess functional marker expression on APC other cells or myeloid other cells, we were able to examine functional marker expression on dendritic cells, CD4^+^ Tregs, and macrophages. We found no significant differences in PD-L1 expression on dendritic cells (Supplementary Figure 5C) or RORGT expression, an immune suppressive functional marker on CD4^+^ Treg cells (Supplementary Figure 5D). Comparison of macrophage functional markers did demonstrate that a significantly greater proportion of macrophages expressed MHCII in KMC tumors versus KPC tumors; while CD206 and PD-L1 expression was consistent across models (Figure 4F). Altogether, these results support the notion that antigen presentation is a distinctive feature of the KMC model.

Unsupervised hierarchical clustering on each of the tissue regions based on the densities of cells present further evaluated differences in TME composition (Figure 4G). Interestingly, while tissue regions somewhat clustered according to genotype, tissue regions sampled from the same mouse (tumor) were often spread across clusters, highlighting the intra-tumoral heterogeneity present in these samples. To quantify this heterogeneity, we calculated the Kullback-Leibler (KL) divergence to assess the difference in immune cell composition of each tissue region compared to the average immune cell composition of the overall tumor from which the region was sampled. Larger KL divergence values indicate more intra-tumoral heterogeneity, while smaller KL divergence values indicate less intra-tumoral heterogeneity, and this metric has been used previously to assess TME heterogeneity.^73,74^ While not statistically significant, we found KMC tumors trended toward having more intra-tumoral heterogeneity in immune cells present as compared to KPC tumors (Figure 4H).

### KMC tumors cluster with PDAc genetic subtypes and exhibit intertumoral heterogeneity

Utilizing bulk RNA sequencing of autochthonous KMC, KM/+C, and KPC tumors, we evaluated transcriptional heterogeneity by principal component analysis (PCA) and sample-to-sample distance analysis on normalized mRNA expression profiles. Tumors formed three distinct clusters (C1, C2, and C3) irrespective of *Myc* zygosity (Figure 5A, and Supplementary Figure 6A). We further investigated the correlation between *Kras* and *Myc* expression, in addition to how this might relate to principal component clustering (Figure 5B). We found a strong correlation between *Kras* and *Myc* expression (Pearson r = 0.74) and a tendency for C3 tumors to be on the lower end of expression and C1 tumors on the higher end, with C2 tumors spanning both groups.

**Figure 5:**
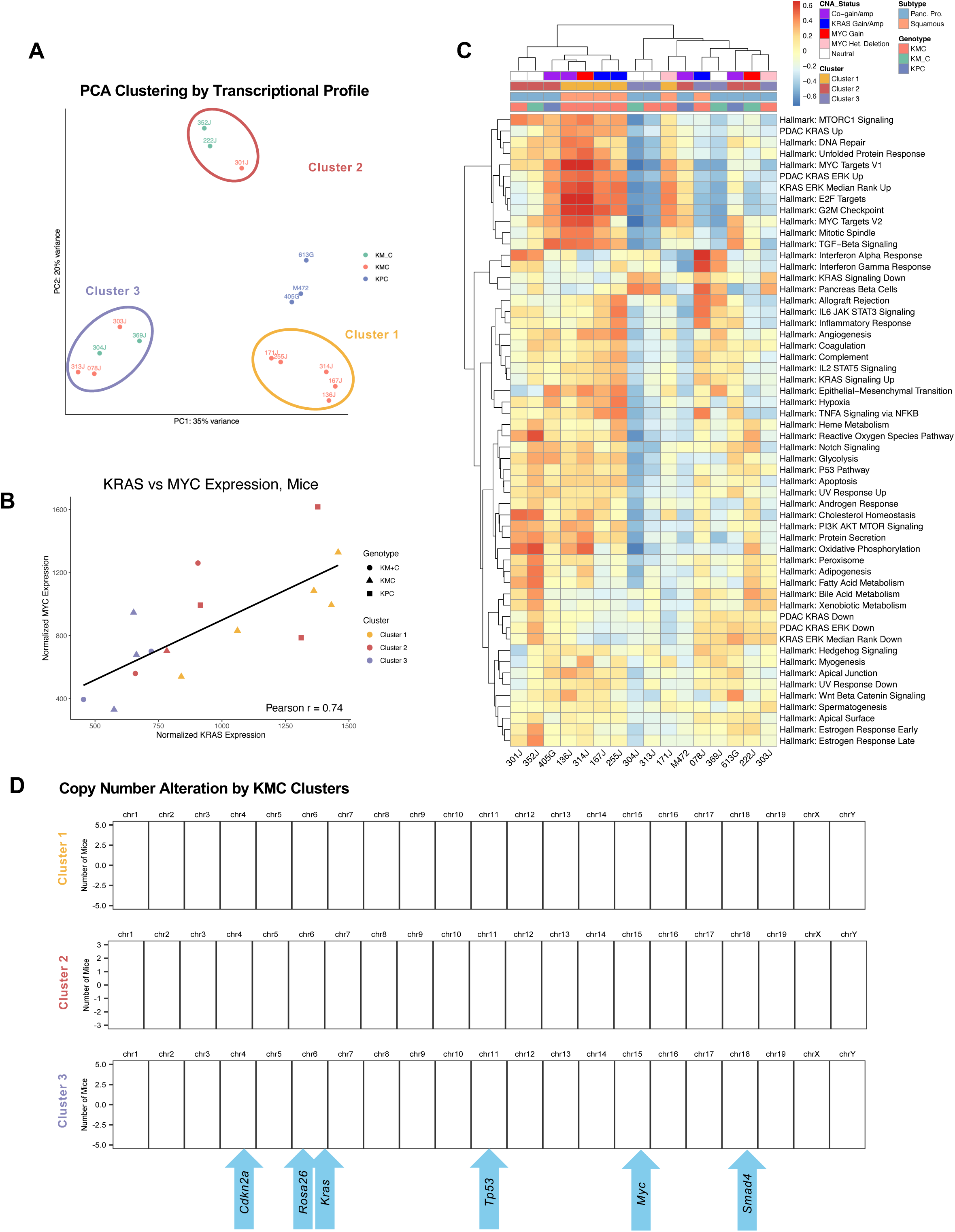
KMC gene expression is characterized by distinctive clustering and transcriptional heterogeneity. A. Principal components analysis of autochthonous KMC, KM/+C, and KPC tumors cluster into three distinctive groups B. Scatter plot with regression line demonstrating correlation between normalized Kras expression and Myc expression in murine tumors (Pearson r correlation coefficient = 0.74), with grouping of transcriptional clustering, irrespective of genotype. C. Heatmap demonstrating sample clustering by GSVA scoring of hallmark gene sets including several curated PDAc Kras specific signatures. Annotation includes copy number alteration for Kras and Myc, transcriptional clustering, PurIST subtyping by shrunken centroid, and model genotype. D. Copy number alteration plots of KMC mice from low pass whole genome sequencing, separated by transcriptional Cluster demonstrating diverse patterns of chromosomal instability in the KMC model; annotated with approximate location of canonical PDAc drivers and KMC alleles.

Tumor transcriptomes were then scored for hallmark gene sets using Gene Set Variation Analysis (GSVA).^75,76^ To evaluate *Kras* mutation signaling, several PDAc RAS-specific signatures were included in the gene set list as well. Clustering on these gene sets yielded differences mainly in *Kras* and *Myc* target gene expression signatures, though distinct groups emerged with respect to inflammatory and infectious responses as well (Figure 5C). Copy-number alterations specific to *Kras* and *Myc* were also evaluated and were identified more commonly in C1 tumors.

We utilized two methods developed to molecularly subtype human PDAc on the KMC and KPC tumor RNAseq data. Purity Independent Subtyping of Tumors (PurIST)^77^, classified all tumors as “Classical”. Since this method is based on eight gene pairs and differences between the human and mouse orthologs could confound the results, we also used the Bailey subtype classification schema and gene signatures that classify Pancreatic Progenitor/Classical versus Squamous/Basal-like.^70^ In this case, tumors classified into each molecular subtype and there was concordance between unsupervised clustering and this subtyping where only squamous tumors were found in C1, while pancreatic progenitor tumors clustered exclusively in C2 and C3 (Figure 5C). The sub-clustering of tumors and alignment with known subtypes of human PDAc supports the ability of this model to recapitulate human disease.

Signatures of genomic structural rearrangement are distinctive to cancer type, with PDAc comprised largely of complex, unclassified rearrangements, deletions, or tandem duplications.^78^ Wanting to evaluate whether transcriptional difference due to clustering were influenced by alterations in copy number we subdivided our low-pass whole genome sequencing into the transcriptomic clusters, identifying distinct alterations across these three groups. C1 was notable for losses in chromosomes 4 (*Cdkn2a*) and 11 (*Tp53*) with gains in chromosome 4 (*Kras* and *Rosa26^Myc^*), and specific gains in *Kras* and *Myc* predominanted in C1 (Figure 5C). C2 had losses in chromosomes 4, 5, and 11 as well, with gains in chromosome 10 & 12. Across C3 there was greater heterogeneity and no clear whole-chromosome ploidy, potentially indicating decreased chromosomal instability or more tumor heterogeneity in this cluster.

Gene Set Enrichment Analysis (GSEA) and the **P**rotein **AN**alysis **TH**rough **E**volutionary **R**elationships (PANTHER) Classification System was applied to contextualize transcriptional differences across tumor clusters.^79–81^. This revealed distinct biological processes between the three KMC tumor clusters. Compared to C2 and C3, we found C1 tumors upregulated key Hallmark gene sets including MYC_V1 and MYC_V2 targets (Supplementary Figure 7) as well as E2F targets, G2M checkpoint, mitotic spindle, epithelial to mesenchymal transition (EMT), TGF-β signaling, and DNA repair. In C2 tumors, we identified upregulation of multiple metabolic processes including oxidative phosphorylation, adipogenesis, glycolysis, and catabolism (Supplementary Figure 8). Immunologic themes predominated in C3 over both C1 and C2, with upregulation of NK cell protection, immune cell chemotaxis, mesenchymal proliferation, and INF-α and INF-γ response pathways identified, in addition to pancreas beta cell and pancreas development gene set enrichment (Supplementary Figure 9). On balance, these data demonstrate the KMC model recapitulates the complex genomic patterns and transcriptomic heterogeneity observed in human disease.

### KMC tumors intercalate with human PDAc representing diverse subtypes and clinical outcomes

To further investigate how the KMC model relates to human disease, we compared tumor RNA sequencing to our center’s large transcriptomic dataset.^24^ After mapping normalized counts of transcripts to human orthologs using Ensembl BioMart^82–84^, counts tables were merged on gene IDs and *ComBat*-seq normalization for batch and species related effects was applied.^85^ Unsupervised clustering by Euclidean distances with Ward.D2 linkage was performed (Figure 6). KMC clusters 1-3 separated into different human PDAc clusters. Metadata for these clusters is shown and includes PurIST classical and basal-like classification^77^ (cf. Moffitt *et al.*^86^) of human samples, single-sample gene-set enrichment analysis of the Hallmark Myc V1 signature and the “pORG” primary PDAc liver organotropism signature described by our center^24^, and clinical metrics including receipt of chemotherapy and survival quartiles from resection to death, last follow up, or dataset censoring.

**Figure 6:**
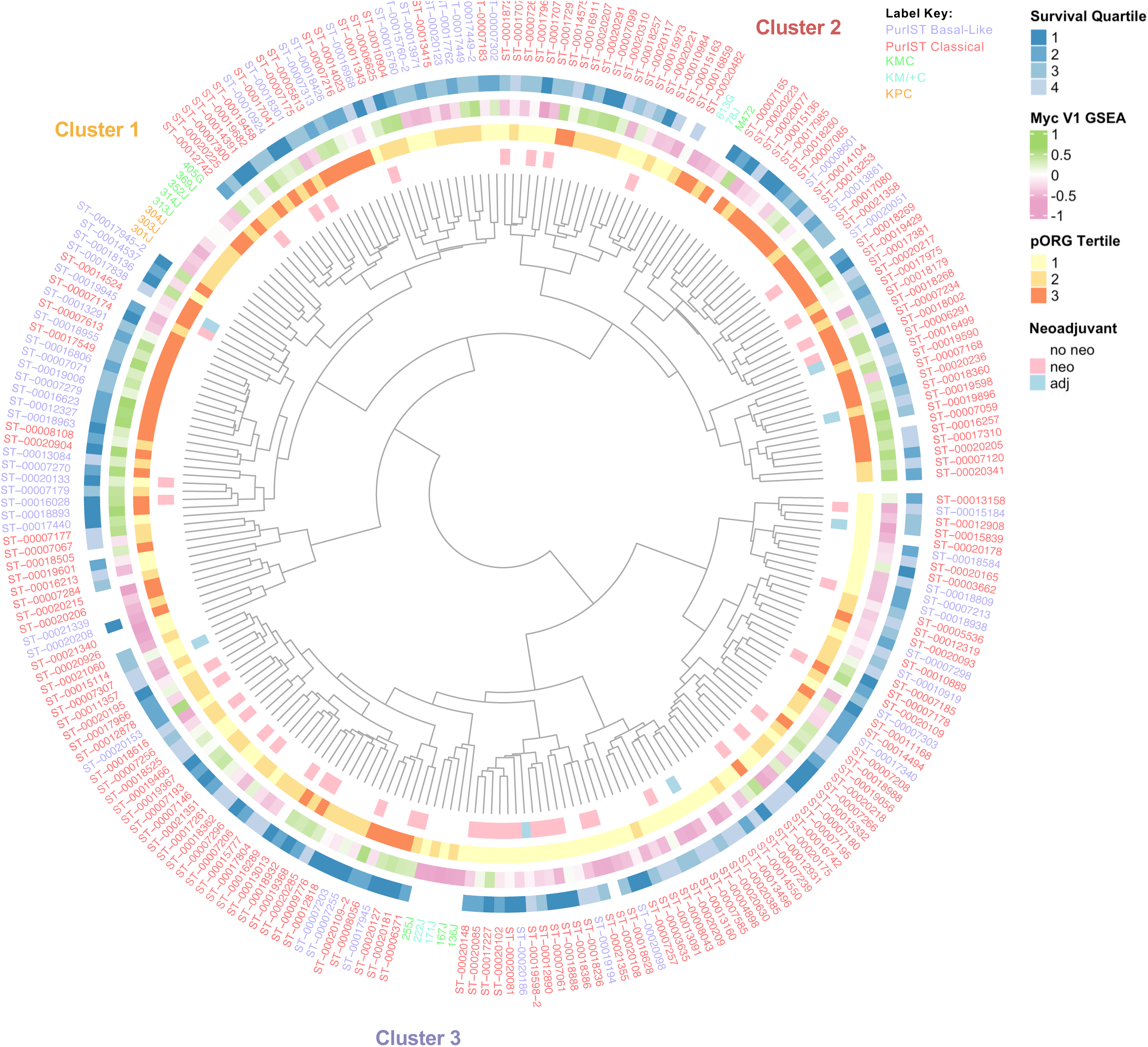
KMC tumors intercalate diversely and by subtype with human PDAc. A. A circular dendrogram of primary human PDAc, KMC, and KPC tumors subjected to RNA sequencing, normalization, homolog mapping, and ComBat normalization. Histogram is by unsupervised Ward D2 clustering. Annotation rings (inner to outermost) describe receipt of chemotherapy (neoadjuvant = pink, adjuvant = blue), pORG score tertile, single-sample GSEA Myc V1 targets score, and survival quartile. Outer ring denotes subject number, colored for humans by PurIST subtyping (blue = basal-like, red = classical) and for mice by genotype (green = KMC, light blue = KM/+C, orange = KPC). The KMC transcriptional clusters (cf. fig 5A) are annotated.

We found that KMC Cluster 1 tumors, which were classified as squamous, fell within a range of majority basal-like human tumors, consistent with the described concordance between these two transcriptional subtypes.^87^ Notably, all three of our sequenced endpoint KPC tumors also fell within this region, which was scored highest in terms of pORG scores as well. Cluster 2 tumors also exhibited high pORG scores and were universally pancreatic progenitor subtype (Figure 5C), consistent with their distribution amongst classical PurIST subtype human tumors. Cluster 3 KMC tumors fell within a region of better survival amongst a higher concentration of patients having received neoadjuvant chemotherapy. These results underscore the range of the KMC system in its ability to model the biological and clinical diversity seen in human PDAc.

### KMC tumor-derived cell lines recapitulate features of autochthonous parent tumors

Due to long latency and impracticality of exclusively relying on an autochthonous model, we generated cell lines from autochthonous tumors (Figure 7A). We established eight KMC tumor cell lines and characterized many of them via multi-omic approaches. Five of the lines underwent WGS and WES (*n* = 4; 171J, 255J, 301J, 303J, and 314J; Supplementary Table 5). When compared to WGS and WES from autochthonous parent tumors, no additional driver alterations were identified (supplemental dataset x). We performed bulk RNA sequencing on seven of the KMC cell lines (171J, 255J, 301J, 303J, 314J, 106L, and 011T). As with autochthonous tumors, cell lines demonstrated heterogeneity in enrichment profiles for Hallmark gene sets (Figure 7B)

**Figure 7:**
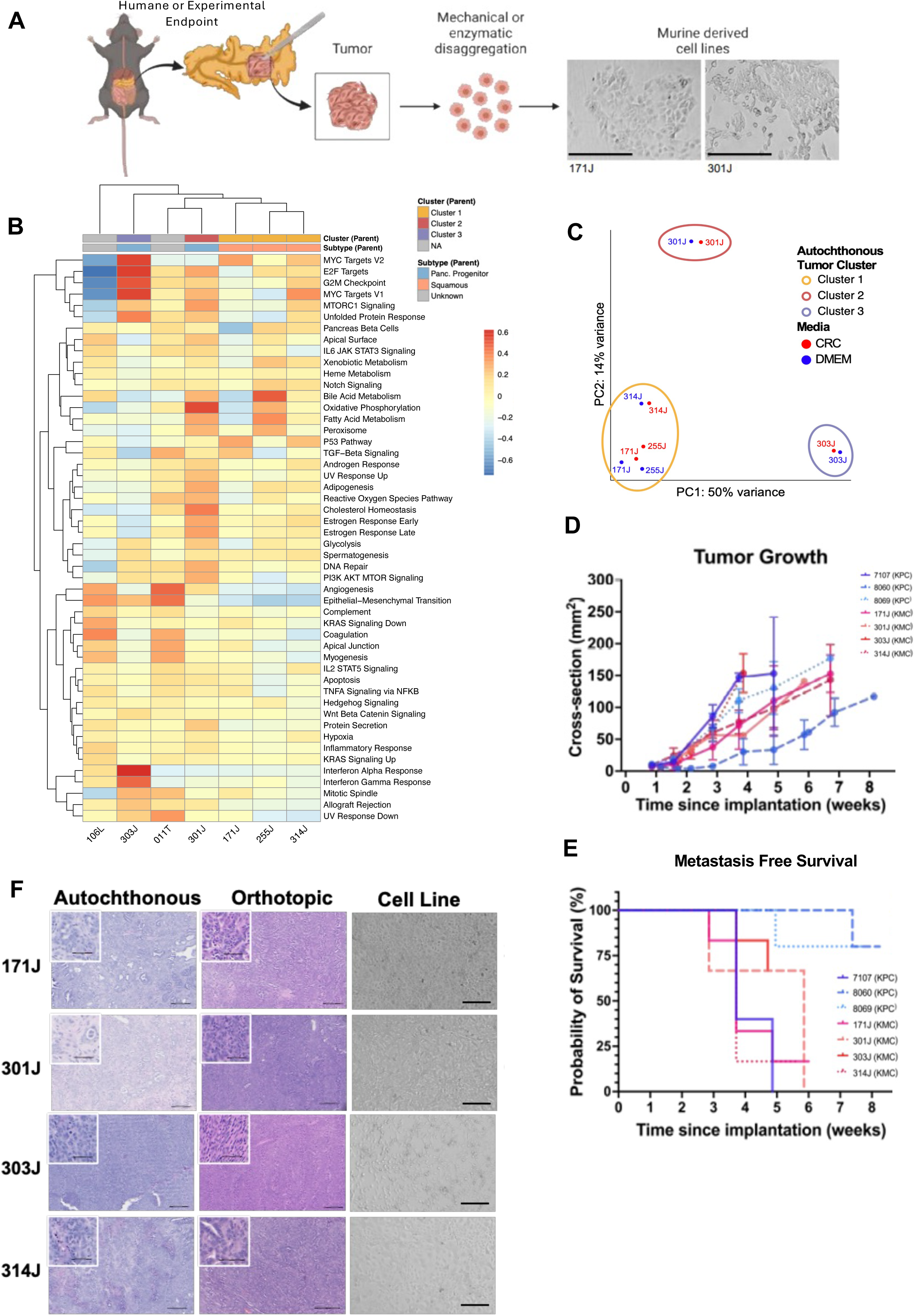
KMC tumor derived cell lines accurately represent parent tumors in terms of histology, growth, clustering, and gene expression. A. Schematic work flow of tumor cell derivation and development of KMC cell lines with representative images of diverse morphology B. Unsupervised clustering of Hallmarks gene expression in cell lines derived from KMC parent tumors with corresponding “clustering” of the parent autochthonous line (cf. fig 5A) and parental tumor PurIST subtyping. C. Principal components plot of cell lines grown from CRC media (generation media) and also DMEM + 10%FBS (maintenance media) demonstrating retained clustering which is also consistent with autochthonous clustering. D. Growth curves (n = 3-5 animals per line) tracked by largest cross-sectional area on transabdominal ultrasound. E. And metastasis free survival by transabdominal ultrasound of animals from the same cohort. F. Representative H&Es showing primary tumor image for autochthonous tumors from which reliable cell lines were derived (left-most column) with sections from orthotopic tumors from cell lines derived from parent tumor, demonstrating persistent morphologic features, heterogeneity, and fibrosis. The right most column depicts the in vitro images of the derived lines. Scale bars 200µm on large images; 50µm on insets.

Lines with deeply-characterized autochthonous parent tumors (171J, 255J, 301J, 303J, 314J) were sequenced across media conditions to demonstrate no major changes in transcriptional profile after culturing in DMEM + 10%FBS versus the CRC establishment media. Principle components analysis was performed and cell lines grouped into three distinct clusters that reflected their respective autochthonous parent tumors, with minimal variation based on media (Figure 7C).

We orthotopically implanted five of the cell lines along with three established KPC cell lines (7107, 8060, 8069) which were included as gold-standard controls into immunocompetent mice to assess *in vivo* growth, metastasis, and fidelity to the parent tumor. Growth was measured over several weeks, twice weekly via trans-abdominal ultrasound to track kinetics and monitor for the development of liver metastases in each line (Figure 7D & E). The 255J line did not consistently give rise to tumors and was excluded from the growth curves. KMC orthotopic tumors gave rise to liver metastases between 4-6 weeks post implantation (Figure 7E and Supplemental Table 5). Analysis of the KMC orthotopic tumors and *in vitro* morphologic features of cell lines showed histologic similarity with the parent tumors from which they were derived (Figure 7F).

### KMC cell lines demonstrate heterogenous treatment response and subtype-dependent chemoresistance

To evaluate treatment response to standard-of-care chemotherapy gemcitabine and abraxane, we selected cell lines of each subtype from two different Clusters: *squamous* (C1, 171J) and *pancreatic progenitor* (C2, 301J) (Figure 8A-D). Abraxane is albumin-bound paclitaxel; as human albumin is immunogenic in mice, paclitaxel alone was substituted. After orthotopic tumors were established, we randomized mice to treatment with standard-of-care gemcitabine 80mg/kg and paclitaxel 15 mg/kg (“GP”) or vehicle via intraperitoneal injection, once weekly dosing. Observation via trans-abdominal ultrasound twice weekly indicated that line 301J appeared to have increased sensistivity to GP. We repeated this experiment with 100 mg/kg and paclitaxel 15 mg/kg (GA) or vehicle I.P., twice weekly and measured clear differences between squamous (171J, Figure 8C) and pancreatic progenitor (301J, Figure 8D) orthotopic tumor growth and overall survival (Figure 8E), with increased sensitivity observed in the 301J line. This finding aligns well with previously reported subtype-dependent drug responses in human tumors: squamous/basal-like tumors were resistant to Gem/Abraxane, and classical/pancreatic progenitor tumors showed a clear response.^88^

**Figure 8:**
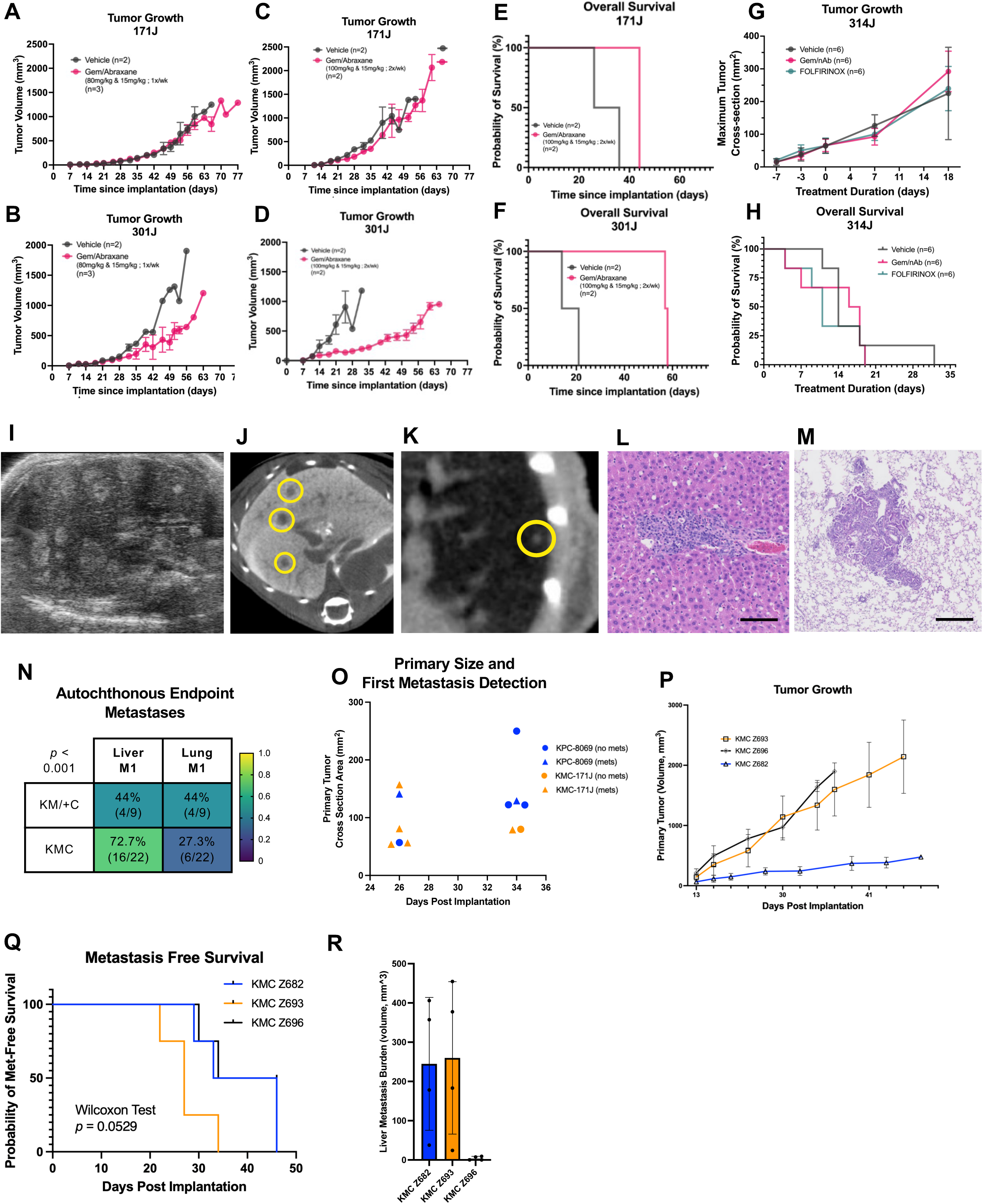
KMC cell lines demonstrate subtype dependent treatment response and reproducible metastatic features with Myc-dose dependent tropism. A. Calculated tumor volume for initial treatment schema with gemcitabine (80mg/kg) and paclitaxel (15mg/kg) once weekly yielded no treatment effect in squamous subtype line 171J, and only modest treatment effect in pancreatic progenitor subtype line 301J B. This treatment response persisted in 171J with C. an increased dosing regimen (gemcitabine 100mg/kg + paclitaxel 15mg/kg twice weekly) in 171J, however increased dose and frequency rendered 301J sensitive D. to treatment. E. Survival analysis detailing no difference in overall survival in 171J with increased gem treatment, but F. increased survival in 301J. G. Subsequent investigations using FOLFIRINOX and gemcitabine/paclitaxel in also confirmed chemoresistance in subsequent treatment in a squamous subtype line (314J) in terms of tumor growth and H. overall survival. I. Representative ultrasound image of hyperechoic liver metastases J. Representative axial liver (left image) and K. lung (right image) MicroCT scans with liver metastases (yellow circles) L. H&E images of a liver metastasis in an autochthonous KMC tumor (left panel; scale bar denotes 50µm) and M. a lung metastasis (right panel; scale bar denotes 200µm) N. Heatmap table of proportion of autochthonous KMC and KM/+C mice with liver and/or lung metastases; p < 0.001 using multiple paired t-tests with two-stage linear setup (cf. Benjamini, Krieger, & Yekutieli) O. Comparison of time of detection of first metastatic disease and tumor size for six KPC (blue) and six KMC (orange) mice. Each dot represents a different mouse. Triangles indicate mets were detected; circles are the final measurement and indicate no mets were detected for that mouse. P. Comparison of mean tumor growth rate and error for BL/6 background KMC lines. Q. Kaplan-Meier curve demonstrating the differential metastatic potential of the BL/6 KMC lines R. Endpoint total metastatic volume by BL/6 KMC cell line.

We wanted to further evaluate the chemosensitivity of lines that were resistant to gem/paclitaxel to FOLFIRINOX. We utilized the squamous-subtype cell lines 171J and 314J and randomized mice to vehicle, gem/Abraxane at the above higher dose, or FOLFIRINOX (see Methods). No differences were observed in either line with respect to tumor growth or overall survival (Figure 8G & 8H; Supplement Fig 10A-C). While additional studies with larger cohort numbers will be required to validate these therapy response results and determine significance, these initial studies demonstrate the potential of KMC-derived lines as a new set of tools with which to study and overcome therapy resistance in PDAc.

### MYC drives a reliably metastatic phenotype with dose-dependent tropism in the KMC model

The majority of patients with pancreatic cancer present with metastatic disease, rendering them ineligible for surgical resection.^89,90^ One of our early observations in the autochthonous KMC model was that gross liver metastases at endpoint were common. On trans-abdominal ultrasound of autochthonous and orthotopic KMC tumor bearing mice, we were able to appreciate progressive, hyperechoic foci that corresponded to metastatic disease in the liver (Figure 8I, Supplementary Fig 10D. This was also observed on contrast enhanced CT scan in the liver (Figure 8J) and the lungs (Figure 8K), a metastatic site that is challenging to image by ultrasound. We confirmed these findings histologically as well in the liver (Figure 8L) and lung (Figure 8M).

MYC is known to affect metastatic patterns in pancreatic cancer.^25^ Our lab has recently shown primary tumor MYC activity to be associated with liver tropism in human pancreatic cancer metastasis.^24^ To investigate whether metastatic patterns varied across KMC genotypes, we reviewed pathologic sections of end-point mice for the presence of liver and/or lung metastases (cf Figures 8L & 8M). Several whole-organ H&E sections from FFPE blocks of lung, liver, and spleen were reviewed for evidence of metastatic disease. In KM/+C tumors, four of nine mice had identified liver metastases and four of nine had lung metastases. KMC homozygous tumors had higher percentage of metastatic disease in the liver (16/22 identified, 72.7%) and a lower percentage had identified lung metastases (6/22, 27.3%, *p <* 0.001, multiple paired *t*-tests). This suggests a dose-responsive tropism, driven by higher Myc activity in the primary tumors, that promotes a liver-avid, lung-averse phenotype (Figure 8N).

We observed consistent development of metastases in the KMC cell lines, with reproducible, line-specific metastatic heterogeneity. Metastases from orthotopic KMC tumors typically appear within 4-6 weeks of implantation and were found to be more consistent than KPC tumors (see above, Figure 7E). These liver metastases were detectable by ultrasound at a smaller primary tumor size in KMC lines when compared to KPC (Figure 8O), thus rendering mice less moribund and allowing for better modeling windows for treatment.

To streamline orthotopic experiments we backcrossed mice with the originating mixed 129/BL6 genotypes which necessitated using 129/BL6 F1 hosts for the orthotopic experiments into pure BL/6 background mice over ten generations and developed additional BL/6 KMC tumor lines. Orthotopic transplant of these lines, Z682, Z693 and Z696 showed differential tumor growth rates tracked via transabdominal ultrasound (Figure 8P) and metastatic potential (Figure 8Q) consistent with lines derived from the mixed-background mice. Tracking endpoint liver metastasis volume in these mice via microCT (Figure 8R) and comparing this with orthotopic growth rate, we observed a complement of differential phenotypes including faster and slower-growing primary tumors with varying timing of metastasis and metastatic burden that can be leveraged to investigate various aspects of PDAc metastasis and biology.

## DISCUSSION

KRAS mutations are ubiquitous and known early events in PDAc initiation, with their presence in both low and high grade PanINs.^10,91^ While *MYC* gene amplification is only documented in ∼10% of primary PDAC, KRAS signaling is known to post-translationally activate MYC through the phosphorylation of Serine 62, which increases MYC protein stability and its DNA binding and oncogenic activity.^18–20,22,30^ This raises the possibility that increased cellular MYC activity through KRAS-mediated stabilization could be an early event in PDAc progression, and forms the basis for developing the KMC model.

Novel to this investigation is the finding that both in humans and in KMC mice, precursor (ADM) and early neoplastic (low-grade PanIN) lesions exhibit elevated levels of stabilized pS62-MYC protein, even absent high levels of gene amplification or mutations, and this increases across disease progression. The consistent development of adenocarcinomas in KMC pancreata —*absent other canonical driver mutations*—underscores a critical role for MYC post-translational regulation by mutant KRAS in PDAc tumorigenesis. Here, we describe the KMC model to further understanding of the progressive development of PDAc and its metastatic dissemination. Time-course sampling of progression in the KMC model demonstrates a dose-dependent development of ADM and PanINs throughout the pancreas that echoes recent work in humans by Braxton & colleagues that unmasked a landscape of genetically-diverse PanINs adjacent to pancreatic tumors which *rarely harbored p53 mutations.* ^10^

Existing pancreatic cancer mouse models leverage known, common oncogene mutations or tumor suppressor loss in human adenocarcinomas, driving these mutations most commonly with embryonic *Pdx1*- or *Ptf1a-Cre*.^92^ While embryonic *Pdx1-Cre*; *LSL*-*Kras^G12D^*^/+^ and *Ptf1a-Cre; LSL-Kras^G12D/+^* mice established the early role of mutant *Kras* in PanIN formation, progression to invasive carcinoma was rare in this model.^93^ Adding mutated LSL-Tp53^R172H/+^ to the Pdx1-Cre version (KPC) of this model accelerated invasive tumor development to ∼3 months and produced frequent metastases, but results in an aggressive model, simultaneously driving early and late mutations, limiting generalizability for the study of early events in tumorigenesis.^44^ Combinations of *Brca2* mutations with KPC mice also yielded aggressive invasive lesions, but models a less common mutation in PDAc.^60,94^ *CDKN2A* mutation and associated loss of p16 (INK4A) was observed in a high proportion of PanINs^95^ and subsequently modelled genetically with a *Pdx1-Cre; LSL-Kras^G12D/+^;LSL-Ink4a^flox/flox^*which yielded aggressive locally invasive tumors and a median survival of two months.^91^ Other models manipulating TGF-β signaling and loss of pathway-intermediary SMAD4—known to be involved in metastasis in human disease—have yielded similar results.^61,96^

We took a different approach by deregulating *Myc* expression at physiologic levels, to facilitate KRAS-mediated post-translationally activation of MYC. This led to tumor evolution which involved the spontaneous suppression of key tumor suppressors p16 and p53, consistent with human disease progression. That driving deregulated *Myc* and mutant *Kras* can spontaneously cause the silencing of p16 or change expression patters of p53 underscores the critical role they play in PDAc development and the utility of the KMC model for early events in tumorigenesis. We continue to investigate the mechanisms of tumor suppressor loss, but one possible explanation is bypass of oncogene induced senescence (OIS), a process that is known to occur in PanINs and other premalignant pancreatic lesions.^97^ OIS results in permanent growth arrest of oncogene driven cells, but accelerates the process of cancer cell evolution by conferring a “senescence activating secretory phenotype”, driving the development of a local microenvironment that is pro-tumorigenic and facilitates malignant transformation.^98,99^

The heterogeneity identified in the KMC model’s tumor microenvironment also is a tool for the study of antigen presentation in PDAc stroma. PDAc is known to have an immunosuppressive microenvironment which is thought to play a role in treatment resistance and poor outcomes.^57,58,100,101^ Dendritic cells are a type of resident tissue antigen presenting cells and play a well-described role in human PDAc.^100^ Dendritic cell paucity is a feature of other canonical PDAc GEMMs, which has necessitated the development of modified or novel systems to model this population.^102–104^ The differences seen in immune densities across phenotype and cluster also underscores the importance of tumor genetics on driving microenvironmental response. This agrees with the finding that antigen presenting cell densities differ across KMC and KPC tumors.

Our observations on the transcriptomic diversity demonstrates utility of the KMC model for studying the spectrum of human PDAc. Autochthonous tumors and the derived cell lines show heterogeneity of molecular subtype including tumors and tumor derived lines representing the two major molecular subtypes: pancreas progenitor/classical and squamous/basal-like. Consistent with published data, we find that KPC tumors cluster entirely with human basal-like tumors^68,105^, whereas the three identified clusters of KMC tumors interpose diversely across our center’s sequenced human PDAc primary tumors, providing biologic variability as well as the possible genetic drivers of subtype determination and clinical outcomes in PDAc.

In addition to gene expression differences, we also identified copy-number variation patterns differed by transcriptional cluster. One interpretation of the heterogeneity in copy number variation across individual clusters within the KMC model--especially as compared to the more homogenous gains/losses observed in our KPC tumors (c.f. Figure 3D bottom row)—is that the latency of the model may allow for greater accumulation of chromothripsis related genomic changes, which has been shown to be present in a majority of human pancreatic cancers.^106^

Treatment resistance and metastatic disease contribute to PDAc lethality: the KMC model offers a novel resource in modeling these aspects of disease. Ways of more effectively testing treatment regimens are also critical to better inform clinical trial design. As with human PDAc, the therapeutic responsiveness of the KMC lines seems related to transcriptional subtype. Recent studies assessing correlations between tumor subtype and patient outcome have shown that squamous PDAc tumors are more aggressive and have worse prognosis.^107–109^ This is observed in the KMC model, particularly with both classical tumors (301J), where susceptibility to gemcitabine/paclitaxel was observed, and squamous/basal-like tumors (171J, 314J) that remain resistant to standard-of-care regimens. That tumors and lines derived from this model share differential subtypes and biology provides a tool for novel experimentation and preclinical modeling. One particular area of interest is the relevance this model will have in testing therapeutics targeting *Kras^G12D^*, given that MYC has been implicated in acquired resistance to RAS-inhibition.^6^

As the principal cause of death in patients with pancreatic cancer, metastasis is one of the most critical features of any PDAc GEMM. KMC cell lines’ reliable metastatic potential, and differential responses to chemotherapy across tumors derived from the KMC model aligns well with the complexity of human disease, providing a robust model for future investigation into metastasis and treatment. Reliable understanding of early events in pancreatic cancer metastasis is difficult in humans given the rarity of early diagnosis and temporal and spatial challenges associated with biopsy. The propensity of PDAc in humans to metastasize even with a small burden of primary disease is also seen in KMC autochthonous and orthotopic tumors, providing an exciting opportunity to evaluate metastatic disease in pancreatic cancer at all stages of dissemination, colonization, and progression. Additionally, differential metastatic potential in this model will provide tools for understanding mechanisms of tropism and pancreatic cancer metastasis.

One resulting limitation of inducing recombination at 5–8-week-old mice is that the latency of tumor development—though universal—is long. We argue that this reflects human disease more accurately but also have derived cell lines that are highly representative of their autochthonous tumor biology and exhibit a reliability and versatility that allows extensive *in vitro* and *in vivo* experimentation.

Orthotopic transplantation has been described as having a “limited” role in modeling of metastasis with other cell lines derived from PDAc GEM models, and frequently relies on intrasplenic injection which results in direct seeding of the portal vein and liver, rather than tumor establishment in the pancreas followed by progression through the metastatic cascade, resulting in a lower-fidelity model of disease.^110^ The predictable metastatic behavior of the KMC lines following orthotopic transplant, as well as the reasonable growth of the primary tumor and metastases allows for tissue collection as well as a treatment window conducive to studying all aspects of pancreatic cancer.

In conclusion, we present here a novel genetically engineered mouse model, and the potential it provides for investigations into critical areas in human PDAc. Confining recombination to the acinar compartment by utilization of *Ptf1a-Cre^ERTM^* with tamoxifen induction at 5-8 weeks of life also abrogated off-target effects seen in embryonic Cre drivers. Resulting tumors develop into adenocarcinomas with high levels of stroma development through a ADM-PanIN-carcinoma progression that can be characterized both histopathologically and with novel machine-learning tissue recognition algorithms. Acquisition of spontaneous p16 and p53 loss of expression further underscores the important role MYC deregulation plays in mediating the early effects of KRAS mutation. Finally, the cell lines derived from this model retain key features from parent tumors including transcriptional subtype, metastatic potential, and *in vivo*/orthotopic behavior, and are already being utilized by collaborators with exciting results.

## MATERIALS AND METHODS

### Primary patient tissue acquisition and patient consent

Primary patient data and tissue were collected through the IRB-approved Oregon Pancreas Tissue Registry (OPTR) (OHSU IRB00003609) in accordance with the Declaration of Helsinki. De-identified tissue was assigned a Participant ID and utilized for relevant experiments (OHSU IRB00003330).

### OHSU Human PDAc Progression TMA

The human progression TMA was designed and generated by the Brenden-Colson Center for Pancreatic Care at Oregon Health & Science University (OHSU). Tissue regions of interest were manually marked by a trained pathologist (T.M.) on FFPE sample blocks from surgically resected primary pancreatic tissue samples from pancreas patients at OHSU. Using a TMA Master II (3DHistech, Hungary) for drilling recipient block and MTA-1 (Estigen Tissue Science, Estonia) for tissue coring, 1.5 mm tissue cores, two cores per surgical specimen (with 4 exceptions where tissue was too small, so only one core was punched), were punched from representative areas containing normal pancreas, pancreatic intraepithelial neoplasia (PanIN, grades 1-3), adenocarcinoma (PDAc, grades 1-3), intraductal papillary mucinous neoplasia (IPMN, grades 1-3), islet cell tumor, and cholangiocarcinoma. Tissue cores were randomized and divided between two paraffin blocks (TMA-1 and TMA-2). The resulting TMAs consisted of 109 unique tissue samples (213 tissue core punches, including duplicates) from 53 patients. Tissue cores included: normal pancreas (n = 17), PanIN grades 1, 2, 3 (n = 10, 7, 1); adenocarcinoma grades 1, 2, 3 (n = 3, 16, 22); IPMN grades 1, 2, 3 (n = 4, 8, 15); islet cell tumor (n = 1); and cholangiocarcinoma (n = 5) samples. There were an additional 6 cores of normal placenta tissue as controls.

### Immunohistochemical staining of OHSU Tissue Micro Array (TMA-1)

TMA-1 was used for the assessment of nuclear pS62-MYC by immunohistochemistry (IHC). The stained section was comprised of tissue samples from 25 patients, including normal pancreas (n = 20), PanIN (n = 7), PDAc (n = 17), and PDAc metastasis (n = 2). Heat-induced antigen retrieval was performed where slides were immersed in target-retrieval solution of citric acid buffer (pH 6.0) for 10 min at 120°C. After antigen retrieval, endogenous peroxide activity was blocked using 3% H_2_O_2_ solution for 15 min at room temperature (RT), followed by protein blocking using 3% FBS/goat serum for 30□min at RT. Next, TMAs were incubated overnight at 4°C primary pS62-MYC antibody (Sears Lab (1:200)). After primary antibodies, TMAs were incubated with anti-rat horseradish peroxidase (HRP) conjugated secondary antibody (Vector Laboratories, cat# BA-9401, 1:150) for 1 hour and ABC Kit (Vector Laboratories, catalog# PK-7100) for 30 min at RT, followed by detection using 3,3′- Diaminobenzidine (Vector Laboratories, catalog# SK-4100) according to manufacturer’s instructions and counterstained using hematoxylin. IHC expression of pS62-MYC was scored as the product of expression level multiplied by percentage of nuclear staining in positive cell.

### Genetically Engineered Mouse Model Generation & Care

All animal studies were conducted in compliance with ethical regulations and were approved by OHSU institutional animal care and use committee (IACUC) (protocol number TR01_IP00001014). The generation and characterization of the Rosa-LSL-Myc mouse (Mus musculus) has been previously described by our lab.^37^ The LSL-Kras^G12D^; Pdx1-Cre mice^93^ were obtained from Dr. Kerry Campbell at Fox Chase Cancer Center. The Ptf1a-Cre^ERTM^ mice have been previously described.^111^ The Tamoxifen-inducible KMC PDAc GEMM was generated by crossing the Rosa-LSL-Myc-HA and LSL-Kras^G12D^ alleles with the conditional Ptf1a-Cre^ERTM^. All mice were on a mixed Bl/6-129 background, and both males and females were used. Allele recombination was induced in mice between 7-9 weeks old with 50 mg/kg Tamoxifen (MilliporeSigma, Burlington, MA) once daily for 5 days by oral gavage.

### Murine Tissue Harvest & Processing

Mice were euthanized when they first appeared moribund or when their pancreatic tumors reached 2 cm^2^ at a maximum tumor cross-section, as measured by ultrasound. Mice in the long-term GEMM survival study not having reached the two aforementioned endpoints at 750 days of age were euthanized at that point. Mice were perfused with 15 ml PBS (Fisher Scientific, Cat. # SH3002802) containing 0.005% heparin (Sigma Aldrich, Cat. # H3393) under general anesthesia. Pancreatic, liver, lung, spleen, gastrocnemius muscle (GM), gonadal fat pad (GFP), and brown adipose (BAT) tissues were harvested and weighed. GM, GFP, BAT and pieces of pancreas, liver, lung, and spleen were preserved in RNAlater and/or flash-frozen in liquid nitrogen. Remaining tissues were then fixed in 10% formalin (4% paraformaldehyde) for 48 hours, then transferred to 70% EtOH. Tissue blocks and slides were made and H&E stained by the Histopathology Shared Resource Core at OHSU or by lab members. Stained slides were scanned using a Zeiss AxioScan microscope at the Advanced Light Microscopy Core at OHSU. Relative pancreas weight was calculated using the following formula:

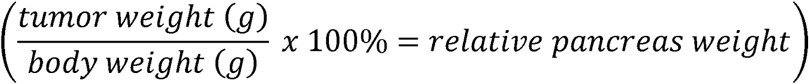

### CODA Tissue Composition Methodology

Mouse pancreas histological composition was quantified using a semantic segmentation deep learning model previously developed for quantification of human pancreas histology.^54^ Manual annotations were performed in 30% of the images to label 10 microanatomical tissue components in pancreatic histology: islets of Langerhans, normal ductal epithelium, pancreatic cancer precursor lesions, pancreatic cancer, vasculature, acinar tissue, fat, collagen, immune hotspots, and non-tissue. A deep learning algorithm was trained to fully label the 10 tissue components in all H&E images to a resolution of 1 μm/pixel. Lymph nodes and duodenum were manually labeled on images where they appeared. Testing annotations were performed in seven histological images and validated by a board-certified pathologist specializing in pancreatic cancer, revealing an independent accuracy of 96% (Supp. Fig. 3A). Following testing, all images were fully segmented by the trained model, and pancreatic compositions were quantified and compared between conditions.

### Immunofluorescence (IF) Tissue Staining

For IF of human samples, formalin-fixed paraffin-embedded tissue sections from patients with matched normal and pancreatic cancer tissues were obtained from archives of the OHSU Department of Pathology, (IRB00002086). These sections were then de-paraffinized, rehydrated, and blocked in 3% BSA. They were incubated in primary antibody (rabbit polyclonal pS62-MYC)^38^ overnight at 4°C. Sections were then incubated with AlexaFluor 488 secondary antibody (Life Technologies, Grand Island, NY) (1:1000) and mounted using SlowFade Gold Antifade mountant with DAPI (Life Technologies, Grand Island, NY). Images were taken with a Hamamatsu digital camera mounted on a fluorescence microscope. Immunofluorescence density was analyzed using the Measure Density tool in OpenLab 5.5 software. pS62-MYC staining intensity was then measured in 10 randomly selected individual cells in each of 5-10 random fields of view and averaged for each sample.

For IF of murine samples, formalin-fixed paraffin-embedded tissue sections were de-paraffinized, rehydrated, and antigen retrieval performed in pH 6 Citrate buffer (Sigma, location) in a pressure cooker for 4 min at high pressure and allowed to de-pressurize for 16 minutes. A second antigen retrieval step was then performed with pH 9 IHC Antigen Retrieval solution (Life Technologies) for 15 min. After rinsing slides in dH2O to cool to room temperature and equilibrating in PBS, tissues were blocked with 10% normal goat serum in PBS for 30 min. Primary antibodies (see Key Resources) were diluted in 5% normal goat serum with 1% BSA in PBS and applied to tissues for overnight incubation at 4°C. Secondary antibodies conjugated to Alexa Fluorophores 488, 555, 568, 594, 647, or 750 were diluted 1:200 in 5% normal goat serum with 1% BSA in PBS and applied to tissues for 1 hr at room temperature. Slides were mounted with SlowFade Gold Antifade mountant with DAPI (Life Technologies).

### Immunohistochemistry (IHC)

Unstained sections were de-paraffinized, rehydrated, and placed in Target Retrieval Solution (Dako, Carpinteria, CA) for 10 min at 100°C. After cooling for 20 min, slides were quenched with 3% hydrogen peroxide for 5 min, followed by blocking with 3% goat serum and primary antibody incubations overnight at 4°C. Primary antibodies used were: CDKN2A (p16-INK4A) (Invitrogen, Waltham, MA), Smad4 (Abcam, Cambridge, UK), p53 (Cell Signaling Technology (CST), Danvers, MA). Sections were then incubated with anti-biotin secondary antibodies (1:1000) and labeling was then detected with the Vectastain ABC kit (Vector Laboratories, Newark, CA). Slides were mounted using Vectamount mounting media (Vector Laboratories).

### IHC Image Processing and Analysis

SMAD4, CDKN2A and P53 IHC single cell staining quantification were performed using color deconvolution in ImageJ followed by the Mesmer algorithm for nuclear segmentation^112^ and a custom algorithm for duct segmentation and signal intensity quantification (https://github.com/engjen/KMC_Mouse). Briefly, CDKN2A and p53 duct segmentation was automated by selecting pixels with DAB and hematoxylin intensity greater than the 0.3 quantile, removing objects smaller than 500 pixels and performing a morphological opening with a 3-pixel disk. SMAD4 duct segmentation was automated by selecting pixels with a hematoxylin intensity greater than the 0.3 quantile, removing holes smaller than 2000 pixels and performing a morphological closing with an 8-pixel disk. ROIs dominated by tumor were manually selected for staining quantification.

### Multiplexed Immunohistochemistry (mIHC), Image Acquisition, and Analysis

Multiplexed IHC was performed on 5 µm formalin-fixed paraffin-embedded (FFPE) sections, as previously described.^101,113–116^ Briefly, slides were deparaffinized and stained with CSF-1R (Santa Cruz Biotechnology, Dallas, TX), F4/80 (Bio-Rad, Hercules, CA), and hematoxylin (Dako, Santa Clara, CA), followed by digital whole-slide scanning at 20X magnification on an Aperio AT2 scanner (Leica Biosystems, Wetzlar, Germany). Tissues then underwent 20 minutes heat-mediated antigen retrieval in pH 6.0 Citra Plus solution (BioGenex, Fremont, CA), followed by endogenous peroxidase blocking in either 0.6% H2O2 for 20 minutes or Dako Dual Endogenous Enzyme Block (Dako, Santa Clara, CA) for 10 minutes. Protein blocking was performed for 10 minutes with 5% normal goat serum and 2.5% BSA in PBS. Slides were incubated with primary antibody for a duration of 30 to 60 minutes at room temperature or overnight at 4°C. Antibodies and staining conditions are listed in Supplementary Table 6. Slides were then washed in TBST and either anti-rat, anti-rabbit, or anti-mouse secondary antibody was applied for 30 minutes at room temperature, followed by signal detection with AEC chromogen (Vector Laboratories, Burlingame, CA). Slides were then digitally scanned before chromogen removal in 100% ethanol. For staining cycles with two rounds of antibody development, protein blocking was repeated after chromogen removal and tissue sections were taken through all steps listed above from primary antibody application through chromogen development, slide scanning, and chromogen removal. The next cycle was then started at the heat-mediated antigen retrieval step, in order to ensure removal of all antibodies from the previous cycle.

### TME Heterogeneity Analyses

*Kullback-Leibler (KL) Divergence*: KL divergence was used to quantify intra-tumoral heterogeneity, as previously published.^73,74^ KL divergence is a statistical measure of how much information is lost when approximating one distribution with a second distribution. Here, we used KL divergence to assess the difference between each tissue region’s distribution of immune cell densities present compared to the average distribution of immune cell densities present for all tissue regions sampled from the same mouse. Larger KL divergence values indicate greater heterogeneity in immune cells present across a given tumor, whereas smaller KL divergence values represent less heterogeneity in immune cells present across the tumor. The entropy function from the scipy Python package (v1.15.2) was used to calculate KL divergence values using a log base of 2.

*Unsupervised Hierarchical Clustering*: Unsupervised hierarchical clustering was used to cluster tissue regions by their cell densities. Densities were log10+1 transformed prior to clustering. The clustermap function from the seaborn Python package (v0.13.2) was used to cluster samples with the method parameter set to ‘average’ and the metric parameter set to ‘cityblock.’

### Tissue Annotation and Region of Interest (ROI) Selection

Whole-slide digitally scanned images for each case were reviewed by a pathologist (TKM), and tumor-enriched tissue regions were digitally annotated. Areas of necrosis were excluded. One FFPE tissue section per GEMM PDAc tumor (n = 3 for KMC cohort; n = 4 for KPC cohort) was used for staining by a combined mouse cell lineage and functional state mIHC biomarker panel. ROI number and size were maintained across samples. To calculate cell densities of immune populations in an individual sample, cell counts for a given cell type (e.g. Dendritic cells) from each ROI were added to get total cell count, and areas of each ROI were also added. Cumulative density was then calculated with the following equation:

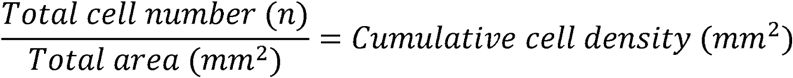

### mIHC Image Processing and Analysis

Image co-registration and processing was performed using methods adapted from the Coussens lab’s previously described image processing workflow.^113,115,117,118^ Briefly, selected ROIs from each RGB marker image were registered to the hematoxylin stained RGB image using the detectSURFfeatures algorithm in the Computer Vision Toolbox in Matlab (version R2018b, The MathWorks, Inc., Natick, MA) to identify matching keypoints in the blue or red channel of the hematoxylin image with keypoints in the blue channel of the AEC stained image. Using estimateGeometricTransform in the Computer Vision Toolbox, a geometric transformation was estimated using similarity that was applied to all channels of the AEC-stained image. This process was used to register each marker-stained image region to the same region of the hematoxylin-stained image.

Image processing, cell quantification, and image cytometry were performed using FIJI (FIJI Is Just ImageJ)^118^, CellProfiler (version 3.5.1)^119^, and FCS Express 6 Image Cytometry RUO (De Novo Software, Glendale, CA), respectively. AEC signal was extracted for quantification and visualization in FIJI using a previously described custom macro [107] for color deconvolution. The resulting data is a csv containing the mean intensity of every cell for each marker. These data are subsequently used in image cytometry single-cell gating, hierarchical identification and quantification in FCS Express (De Novo Software). Hierarchical cell classifications are shown in Supplementary Table 3. For mIHC visualization, signal-extracted images were overlaid in pseudo-color in FIJI.^118^

### MYC genomic alteration survival analysis in published human patient cohort

cBioPortal for Cancer Genomics (http://cbioportal.org) provides a framework for multidimensional cancer genomics data analysis and visualization.^120,121^ In this study, we performed survival analyses based on mutations and copy number alterations (amplifications) in c-MYC appearing in dataset downloaded from cBioPortal: The Pancreatic Adenocarcinoma (TCGA, PanCancer Atlas).

### DNA and RNA isolation

DNA and RNA were isolated from FFPE end-stage murine pancreatic tumor tissue by first identifying tumor-dominant ROIs on full tissue scans of H&E slides for each tumor. These ROIs were used as reference microdissection maps while carefully scraping tumor-rich regions from freshly cut 10 μm sections from slides under a dissection microscope. RNA was then isolated from the microdissected FFPE tissue using an AllPrep® DNA/RNA FFPE kit (Cat. # 80234) with on-column DNase treatment (Qiagen, Hilden, Germany), per manufacturer’s protocol.

KMC cell lines were cultured to 80% confluence on 10 cm cell culture dishes in CRC media or DMEM + 10% FBS. DNA and RNA were isolated directly from 80% confluent 10 cm cell culture dishes for each cell line, using an AllPrep® DNA/RNA mini kit (Cat. # 80204) with on-column DNase treatment (Qiagen, Hilden, Germany), per manufacturer’s protocol.

### Whole Genome Sequencing (WGS) for Copy Number Variant Calling

Library preparation and sequencing were done by the OHSU Massively Parallel Sequencing Shared Resource (RRID SCR_009984). Sequencing libraries for FFPE GEMM tumors were constructed using the Illumina TruSeq Nano Library Prep kit (Illumina, San Diego, CA). Sample libraries were diluted and applied to an Illumina NovaSeq 6000 (RTA version 3.4.4) flow cell at a concentration appropriate to generate about 400 million reads. All libraries were prepared with indexing barcodes to permit multiplexing in a single lane. The 150-cycle paired-end read sequencing was used to assemble the reads into standard FastQ formatted data.

Low-pass whole genome sequencing was performed on tumor/normal pairs first by aligning short-read sequencing data with BWA mem with default settings.^122,123^ Second, ichorCNA (https://github.com/broadinstitute/ichorCNA, Broad Institute, Cambridge, MA)^124^ was used to predict CNVs and estimate tumor fraction with default settings.

### Whole Exome Sequencing (WES) for Somatic Point Mutation Calling

Library preparation and sequencing were done by CD Genomics (Shirley, NY). Briefly, sequencing libraries were generated using SureSelectXT Mouse All Exon Library Prep Kit following manufacturer’s recommendations (Agilent Technologies, Folsom, CA) to enrich genomic DNA for coding exons. Index codes were added to attribute sequences to each sample. Sample libraries were then subjected to paired-end whole exome sequencing using Illumina SBS instruments with a raw sequencing coverage of 200 x. Across samples, the average sequencing depth was 1.28 x 1010 base pairs sequenced per sample.

The sequenced reads were aligned to the mouse genome with BWA mem with default settings.^122,123^ Tumor/normal mutation calling was performed with the NF-Core/Sarek (https://nf-co.re/sarek)^125,126^ pipeline for detection of somatic variants with exome data. The resulting mutation calls were filtered using the Filters for Next Generation Sequencing (FiNGS) package (https://github.com/cpwardell/FiNGS)^127^ using standard settings.

### RNA Sequencing (RNAseq)

Library preparation and sequencing were done by the OHSU Massively Parallel Sequencing Shared Resource (RRID SCR_009984).

Sequencing libraries for FFPE GEMM tumors and KMC cell lines were constructed using the Stranded RNA Prep, Ligation, with the Illumina Ribo-Zero Plus kit or the Illumina TruSeq Stranded mRNA Library Prep kit (Illumina, San Diego, CA), respectively. Briefly, poly(A)+ RNA was isolated from 400 ng of total RNA (per sample) using oligo-dT-coated magnetic beads. The recovered RNA was then chemically fragmented. First-strand cDNA was generated using random hexamers as primers for reverse transcriptase. Strand-specific second-strand cDNA synthesis was performed and a single ‘A’ nucleotide was then added to each end. Illumina adaptors were ligated to the cDNAs. Limited-round (15-cycle) PCR was used to amplify the material to yield the final libraries. Library concentration was determined using real-time PCR with primers complementary to the Illumina adaptors. Sample libraries were diluted and applied to an Illumina NovaSeq 6000 (RTA version 3.4.4) flow cell at a concentration appropriate to generate about 400 million reads. All libraries were prepared with indexing barcodes to permit multiplexing in a single lane. The 100-cycle paired-end read sequencing was used to assemble the reads into standard FastQ formatted data.

RNA-seq analysis was performed similar to previous description.^23^ Briefly, paired-end FastQ sequences were trimmed using trim-galore (version 0.6.3) and default parameters. Pseudoalignment was performed with kallisto (version 0.44.0)^128^ using genome assembly GRCm39 (release M26) and gencode (version 24) annotation; default parameters were used other than the number of threads. The Bioconda package bioconductor-tximport (version 1.12.1) was used to create gene level counts and abundances (TPMs). Quality checks were assessed with FastQC (version 0.11.8) and MultiQC (version 0.1.7). Quality checks, read trimming, pseudoalignment, and quantitation were performed via a reproducible snakemake pipeline, and all dependencies for these steps were deployed within the anaconda package management system.^128,129^

### Differential Expression Analysis

All differential expression analysis was performed in R (version 4.1.0)^130^ using the package DESeq2 (version 1.34.0).^131^ To compare KMC mice to KPC mice, we first summarized Kallisto transcript abundance outputs for all mice into a gene-level expression dataset via tximport (version 1.22.0)^132^, then ran DESeqDataSetFromTximport() to construct the DESeq object used for downstream analysis. Genes with less than 10 cumulative read counts summed across samples in the DESeq object were excluded. Differential expression was performed using the DESeq() function set to compare KMC mice with KPC mice (KMC and KM/+C mice were regarded as a single group). Volcano plots of differential protein-coding genes were created with ggplot2 (version 3.4.1).^133^ For heatmap visualization, we applied variance-stabilizing transformation (VST) to expression counts using DESeq2 and plotted the transformed counts with pheatmap (version 1.0.12).^134^ Heatmap values were scaled by row/gene, and only protein coding genes called differential by an adjusted p-value of 0.05 or less were included. For differential expression between KMC and KM/+C samples, as well as differential expression between KMC & KM/+C clusters, we performed the same computational workflow but excluded KPC mice from the DESeq object and model.

### Analysis of KMC expression clusters

To identify distinct clusters within the KMC PDAc model, we analyzed RNA-seq profiles of homozygous and heterozygous ROSA-MYC-expressing mice (KMC and KM/+C). We applied the R function `dist()` to a matrix of VST-transformed expression counts generated from the KMC & KM/+C DESeq object. Sample-to-sample distances generated through `dist()` were visualized and clustered with pheatmap. Hierarchical clustering results from pheatmap indicated three major clusters that we used for further analysis. To perform PCA, we applied the DESeq2 function `plotPCA()` on the same VST-transformed count data used for clustering. PCA settings in DESeq2 was set to use the top 500 variable genes. Differential expression between clusters was tested in pairwise fashion via DESeq2. Protein-coding genes called significant (adjusted *p*-value of 0.05 or less) within each differential expression test were collected and visualized as a heatmap via pheatmap. VST- transformed counts were used as input for `pheatmap()` and heatmap values were scaled by row/gene.

### Gene ontology (GO) enrichment

All GO^80,135,136^ enrichment analysis was done using the PANTHER^81,137^ statistical overrepresentation test provided by their website (http://www.pantherdb.org). Protein coding genes identified as differentially expressed (min. adjusted p-value 0.05 and abs. log2-fold change 1.0) were used as the test set while the reference set included all protein coding genes with at least 10 read counts summed across samples. PANTHER overrepresentation test settings used were: Fisher’s Exact as the test type, FDR for p-value correction, and “GO biological process complete” as the annotation data set (GO annotation release version 2023-03-06).

### Gene set variation analysis (GSVA)

All GSVA was performed with the GSVA R package (v.1.42.0)^75^ on MSigDB^79^ hallmark gene sets imported with msigdbr (v.7.5.1).^138^ To assign GSVA scores to each sample, VST-transformed counts obtained from DESeq2 were inputted into the gsva() function using default settings. VST-transformation and GSVA were performed on mouse and human RNA-seq datasets separately. Heatmap visualization of GSVA results for KMC & KM/+C samples was done with pheatmap. To improve visual interpretability, dendrogram nodes for samples/columns were rotated to keep clusters 1-3 together without affecting meaning.

### Gene set enrichment analysis (GSEA)

GSEA tool^79,139^ was used with default settings (1000 permutations) on RNAseq clusters data to compare enrichment to gene ontology and hallmark gene sets in the MSigDB database (version 7.5.1) (see Supplementary dataset 3).^140^

### Survival analysis of human pancreatic cancer RNA-seq profiles

To analyze the impact of MYC on overall survival in PDAc, we used existing human pancreatic tumor RNA-seq profiles obtained from the Oregon Pancreas Tissue Registry under IRB study # IRB00003609. Informed consent was obtained from all subjects and experimental protocols were approved by the OHSU Institutional Review Board. The collection, processing, and sequencing of these samples are described by Link JM, et al.^24^ Clinical information pertaining to specimen site, time from diagnosis to death or follow-up, and vital status at last follow-up was collected from the OHSU Cancer Registrar on 01-26-2023. Samples were included for the present analysis if the sequenced specimen was obtained from a primary tumor and the clinical annotation specifying tumor site included “Pancreas”. RNA-seq profiles (formatted as Kallisto transcript abundance files) for each sample were imported into a DESeq object through the same general tximport workflow we used for the mouse differential expression analysis. To perform Kaplan-Meier survival estimation, we used the R package survival (version 3.3-1)^141^ and plotted the resulting survival curves with survminer (version 0.4.9)^142^

### Primary Murine Cell Line Generation

Cell lines 011T, 106L, 171J, 255J, 301J, 303J, 314J, and 808J were propagated from primary murine PDAc tumor tissue and cultured in conditionally reprogrammed cell (CRC) media (Ham’s F-12:DMEM (1:3) media supplemented with 5% FBS, 0.4 μg/mL hydrocortisone, 5 μg/mL human insulin, 8.4 ng/mL cholera toxin, 10 ng/mL EGF, 24 μg/mL Adenine, 10 μM ROCK inhibitor Y-27632, 0.25 μg/mL Amphotericin B, and 50 mg/mL Primocin).^143^ Tumors were collected sterilely, rinsed in PBS, minced with sterile scalpels and further mechanically dissociated in CRC media using a Medimachine instrument (BD Biosciences, Franklin Lakes, NJ). Resulting suspension was then transferred to a 6- well dish and incubated for 24-48 hrs before further manipulation. Serial trypsinization with 0.05% Trypsin was performed every 3 days or when plate reached 90% confluency in order to remove fibroblasts and select for epithelial cancer cell propagation. Serial trypsinization was stopped when exclusively epithelial cancer cells were observed. Cell lines were considered established when doubling efficiently after 3 passages. Once established, cell lines were confirmed mycoplasma-free and used between passages 5-10 for orthotopic experiments and before passage 15 for *in vitro* experiments. 2D cultures were maintained in CRC media unless otherwise noted.

### Orthotopic Allograft Model Generation and Methodology

Pancreatic orthotopic allografts were performed as previously described.^144^ Briefly, 8-12 week-old B6129SF1/J mice (Cat. # 101043, The Jackson Laboratory, Bar Harbor, ME) were given meloxicam as an analgesic and then anesthetized with isoflurane. A small abdominal flank incision was then made, and the spleen exteriorized. Cells were resuspended in 50 μL CRC media and Matrigel (1:1) and 25 μL were injected into the tail of the pancreas using a 28-gauge needle (injection numbers and *n* for each experiment in text). After injection, the needle was held in place for 30 seconds to allow for partial solidification of cell suspension, and the injection site was compressed with a sterile cotton swab as the needle was removed. All exteriorized tissue was then placed back into the body cavity, the abdominal wall and peritoneum sutured closed with 5-0 vicryl, and the skin closed with Vetbond tissue adhesive (3M, Saint Paul, MN). Animals were monitored daily for 1 week post-injection for overall health and wound healing.

### *In Vivo* Drug Studies

After the tumors reach a size of ∼60 mm2 (∼2 wks), mice were divided into cohorts based on average tumor size and rate of tumor growth. All therapeutic agents were obtained from SelleckChem (Houston, TX) except for Gemcitabine (LC Laboratories, Woburn, MA). For gemcitabine/paclitaxel treatments^145,146^, mice were treated IP with 50 mg/kg Gemcitabine (LC Laboratories) and 0.5 mg/kg paclitaxel once weekly. For FOLFIRNOX treatments^147^, mice were treated IP with 100 mg/kg folinic acid (FOL, also known as leucovorin) + 5 mg/kg 5-fluorouracil (F) + 50 mg/kg irinotecan (IRIN) + oxaliplatin 5 mg/kg (OX) in 10% DMSO on days 14 and 16. Note: typical administration of these drugs is performed in 30-day cycles; a second cycle of treatment would be administered on days 44 and 46.

### Tumor Size and Metastatic Disease Assessment

Tumor size and presence of metastases were monitored over time by ultrasound using a Vevo-LAZR 2100 high-frequency/high-resolution linear array Ultrasound (FUJIFILM VisualSonics, Toronto, CA). Upon detection, tumors were imaged and measured using the Vevo 770 image system software. Tumors were continuously monitored by ultrasound over time (weekly for long-term survival study; biweekly for orthotopic allograft studies).

### Western Blotting

Cells were lysed in AB Lysis buffer and Western blot analysis was performed. Briefly, equal amounts of protein were fractionated on 4% to 12% Tris-glycine polyacrylamide gels (Criterion gels, Bio-Rad, Hercules, CA), transferred to polyvinylidene difluoride membranes (PVDF), and probed with primary antibodies against CDKN2A (p16INK4A (E5F3Y), 1:1000), SMAD4 (D3R4N, 1:1000), P53 (D2H9O, 1:1000), and GAPDH (6C5; 1:5000). Antibodies were diluted in blocking buffer (1:1 Odyssey Blocking Buffer (LI-COR Biosciences, Lincoln, NE): PBS with 0.001% Tween20). Primary antibodies were detected with secondary antibodies (1:10,000) labeled with the near-infrared fluorescent dyes IRDye800 (Rockland, Philadelphia, PA) and Alexa Fluor 680 (Molecular Probes, Eugene, OR) diluted in blocking buffer. Blots were visualized and bands quantified on an Odyssey imaging system (LI-COR Biosciences).

### Software

R (version 4.2.3) was used with R packages DESeq2, GSVA, msigdbr, and ggplot. GSEA was run using GSEA tool (version 4.3.2).^79,139^ Western blots were quantified and analyzed using Image Lab software (version 6.1.0, Bio-Rad, Hercules, CA).

## Statistical Methods

All statistical tests were performed with GraphPad Prism 9 or R (Version 4.2.3). A Mantel-Cox test was used to compare all Kaplan-Meier survival and recurrence curves. ANOVA was used for plots with multiple comparisons and t-tests were used for plots with single comparisons.

## Supplementary Information

Supplementary Dataset 1: RNA-sequencing TPM values for all genes

Supplementary Dataset 2: DESeq2 analysis of RNA-seq

Supplementary Dataset 3: GSEA analysis of RNA-seq

Supplementary Dataset 4: WGS Raw FastQ files

Supplementary Dataset 5: CNA analysis of WGS

Supplementary Dataset 6: WES Raw FastQ files

Supplementary Dataset 7: SNV analysis of WES

The datasets generated during this study will be available at the NCBI Gene Expression Omnibus; dbGap submission and accession codes in progress.

## Code Availability

All code is generated through open-source tools and available at https://github.com/patrickjworth/kmc. A README.txt in this repository includes locations of source datasets.

**Supplementary Figure 1:** Embryonic Ptf1a-Cre driving KRASG12D and deregulated MYC results in an aggressive model of diverse pancreatic neuroendocrine carcinomas, neoplasia, and adenocarcinomas

A. Schema of the embryonic (noninducible) KMC Ptf1a-Cre genotype

B. Kaplan-Meier survival curve demonstrating overall survival of non-inducible Ptf1a-Cre driven model showing shorter latency of the model but also that Myc deregulation alone is insufficient to drive PDAc formation.

C. Representative images by genotype of time-point mice aged to two-, four-, six-, and twelve-months. Note the early development of gland-forming neoplasia at two months in this model.

**Supplementary Figure 2:** Kaplan-Meier curve of KMC-model genotypes with number at risk.

A. Cancer specific survival by genotype with number at risk for KMC model genotypes

**Table 1**: Characteristics of autochthonous KMC and KM/+C mice subjected to RNA sequencing including major assays and derived cell lines.

**Supplementary Figure 3:** Supplementary materials for CODA histotyping

A. Confusion matrix demonstrating validation of machine learning algorithm for tissue segmentation.

Diagonal indicates “correct” calls from the algorithm based on pathologist-defined “ground truth”.

B. Comparisons of tissue breakdown based on CODA calls by genotype.

**Table 2**: Results from low-pass whole genome sequencing on autochthonous KMC tumors

**Supplementary Figure 4:** Heatmap demonstrating copy number alteration analysis by tumor with unsupervised clustering.

**Table 3**: Antibody details and multiplex immunohistochemistry details for TME characterization of endpoint tumors.

**Table 4**: Gating strategy for staining results from mIHC to call specific cell types.

**Supplementary Figure 5:** Supplementary figures on transcriptional and immune microenvironmental heterogeneity

A. Volcano plot of differential expression results evaluating transcriptional differences by GEMM model (KPC vs KMC/KM+C); Significance set at p ≤ 0.05; |log2 fold-change| > 1

B. Box plot showing total immune and neoplastic cell densities split by genotype. Each dot represents one tissue region (n= 18 KMC, n= 22 KPC). (p = NS)

C. Box plot showing the percent of dendritic cells positive for PD-L1 functional marker split by genotype. Each dot represents one tissue region (n=18 KMC, n= 12 KPC). Tissue regions with fewer than 15 dendritic cells present were removed from this analysis. (p = NS)

D. Box plot showing the percent of CD4+ Tregs positive for RORGT functional marker split by genotype. Each dot represents one tissue region (n=15 KMC, n= 14 KPC). Tissue regions with fewer than 15 CD4+ Tregs present were removed from this analysis. (p = NS)

**Supplementary Figure 6:** Supplementary figures detailing transcriptomic clustering

A. Confusion matrix demonstrating the generation of three clustering groups from transcriptomic data.

B. Representative H&E images of phenotypic heterogeneity seen among KMC tumors by cluster.

Scale bars depict 200 m.

**Supplementary Figure 7:** KMC Cluster 1 tumors are upregulated in cell cycle, DNA replication and repair biological process GO Terms and GSEA hallmark pathways.

A. Significantly upregulated GO ontology terms for Biological Processes in C1 vs C2

B. Significantly upregulated GO ontology terms for Biological Processes in C1 vs C3

C. Gene set enrichment plots demonstrating significantly upregulated Hallmark gene sets comparing C1 vs C2+C3

**Supplementary Figure 8:** KMC Cluster 2 tumors are upregulated in oxidative phosphorylation and metabolic biological process GO Terms and GSEA hallmark pathways.

A. Significantly upregulated GO ontology terms for Biological Processes in C2 vs C1

B. Significantly upregulated GO ontology terms for Biological Processes in C2 vs C3

C. Gene set enrichment plots demonstrating significantly upregulated Hallmark gene sets comparing C2 vs C1+C3

**Supplementary Figure 9:** KMC Cluster 3 tumors are upregulated in immune-modulatory and tyrosine kinase biological process GO Terms and GSEA hallmark pathways.

A. Significantly upregulated GO ontology terms for Biological Processes in C3 vs C1

A. Significantly upregulated GO ontology terms for Biological Processes in C3 vs C2

A. Gene set enrichment plots demonstrating significantly upregulated Hallmark gene sets comparing C1 vs C2+C3

**Table 5**: In vivo and in vitro characteristics of cell lines generated from KMC parent lines.

**Supplementary Figure 10:** Additional treatment data for standard-of-care chemotherapies in a squamous-subtype line (171J) showing A. overall survival, B. tumor growth and C. time to metastasis.

Representative ultrasound images of liver metastases in a mouse with an orthotopic tumor.

## References

1 Cancer statistics, 2024 - Siegel - 2024 - CA: A Cancer Journal for Clinicians - Wiley Online Library.

2 Cancer statistics, 2023 - Siegel - 2023 - CA: A Cancer Journal for Clinicians - Wiley Online Library.

3 Conroy, T. et al. FOLFIRINOX versus Gemcitabine for Metastatic Pancreatic Cancer. N Engl J Med 364, 1817–1825 (2011). 10.1056/NEJMoa1011923

4 Conroy, T. et al. FOLFIRINOX or Gemcitabine as Adjuvant Therapy for Pancreatic Cancer. N Engl J Med 379, 2395–2406 (2018). 10.1056/NEJMoa1809775

5 Awad, M. M. et al. Acquired Resistance to KRAS(G12C) Inhibition in Cancer. N Engl J Med 384, 2382–2393 (2021). 10.1056/NEJMoa2105281

6 Dilly, J. et al. Mechanisms of Resistance to Oncogenic KRAS Inhibition in Pancreatic Cancer. Cancer Discov 14, 2135–2161 (2024). 10.1158/2159-8290.CD-24-0177

7 Witkiewicz, A. K. et al. Whole-exome sequencing of pancreatic cancer defines genetic diversity and therapeutic targets. Nat Commun 6, 6744 (2015). 10.1038/ncomms7744

8 Mendiratta, G. et al. Cancer gene mutation frequencies for the U.S. population. Nat Commun 12, 5961 (2021). 10.1038/s41467-021-26213-y

9 Sinkala, M. Mutational landscape of cancer-driver genes across human cancers. Sci Rep 13, 12742 (2023). 10.1038/s41598-023-39608-2

10 Braxton, A. M. et al. 3D genomic mapping reveals multifocality of human pancreatic precancers. Nature, 1–9 (2024). 10.1038/s41586-024-07359-3

11 Habbe, N. et al. Spontaneous induction of murine pancreatic intraepithelial neoplasia (mPanIN) by acinar cell targeting of oncogenic Kras in adult mice. Proc. Natl. Acad. Sci. U.S.A. 105, 18913–18918 (2008). 10.1073/pnas.0810097105

12 Risques, R. A. & Kennedy, S. R. Aging and the rise of somatic cancer-associated mutations in normal tissues. PLOS Genetics 14, e1007108 (2018). 10.1371/journal.pgen.1007108

13 Fiala, C. & Diamandis, E. P. Mutations in normal tissues—some diagnostic and clinical implications. BMC Medicine 18, 283 (2020). 10.1186/s12916-020-01763-y

14 Maurice, D. et al. Loss of Smad4 Function in Pancreatic Tumors. Journal of Biological Chemistry 276, 43175–43181 (2001). 10.1074/jbc.M105895200

15 Jones, S. et al. Core Signaling Pathways in Human Pancreatic Cancers Revealed by Global Genomic Analyses. Science 321, 1801–1806 (2008). 10.1126/science.1164368

16 Shain, A. H., Salari, K., Giacomini, C. P. & Pollack, J. R. Integrative genomic and functional profiling of the pancreatic cancer genome. BMC Genomics 14, 624 (2013). 10.1186/1471-2164-14-624

17 Raphael, B. J. et al. Integrated Genomic Characterization of Pancreatic Ductal Adenocarcinoma. Cancer Cell 32, 185–203.e113 (2017). 10.1016/j.ccell.2017.07.007

18 Sears, R., Leone, G., DeGregori, J. & Nevins, J. R. Ras Enhances Myc Protein Stability. Molecular Cell 3, 169–179 (1999). 10.1016/S1097-2765(00)80308-1

19 Sears, R. et al. Multiple Ras-dependent phosphorylation pathways regulate Myc protein stability. Genes Dev 14, 2501–2514 (2000). 10.1101/gad.836800

20 Sears, R. C. The life cycle of C-myc: from synthesis to degradation. Cell Cycle 3, 1133–1137 (2004).

21 Yeh, E. et al. A signalling pathway controlling c-Myc degradation that impacts oncogenic transformation of human cells. Nat Cell Biol 6, 308–318 (2004). 10.1038/ncb1110

22 Farrell, A. S. et al. MYC regulates ductal-neuroendocrine lineage plasticity in pancreatic ductal adenocarcinoma associated with poor outcome and chemoresistance. Nat Commun 8, 1728 (2017). 10.1038/s41467-017-01967-6

23 Langer, E. et al. The Prolyl Isomerase PIN1 Plays a Critical Role in Fibroblast Differentiation States to Support Pancreatic Cancer. SSRN Journal (2021). 10.2139/ssrn.3800381

24 Link, J. M. et al. Ongoing replication stress tolerance and clonal T cell responses distinguish liver and lung recurrence and outcomes in pancreatic cancer. Nat Cancer 6, 123–144 (2025). 10.1038/s43018-024-00881-3

25 Maddipati, R. et al. MYC Levels Regulate Metastatic Heterogeneity in Pancreatic Adenocarcinoma. Cancer Discovery 12, 542–561 (2022). 10.1158/2159-8290.CD-20-1826

26 Gabay, M., Li, Y. & Felsher, D. W. MYC Activation Is a Hallmark of Cancer Initiation and Maintenance. Cold Spring Harb Perspect Med 4, a014241 (2014). 10.1101/cshperspect.a014241

27 Hessmann, E., Schneider, G., Ellenrieder, V. & Siveke, J. T. MYC in pancreatic cancer: novel mechanistic insights and their translation into therapeutic strategies. Oncogene 35, 1609–1618 (2016). 10.1038/onc.2015.216

28 Ni, C. et al. PI3K/ c-Myc/AFF4 axis promotes pancreatic tumorigenesis through fueling nucleotide metabolism. Int. J. Biol. Sci. 19, 1968–1982 (2023). 10.7150/ijbs.77150

29 Schleger, C., Verbeke, C., Hildenbrand, R., Zentgraf, H. & Bleyl, U. c-MYC Activation in Primary and Metastatic Ductal Adenocarcinoma of the Pancreas: Incidence, Mechanisms, and Clinical Significance. Modern Pathology 15, 462–469 (2002). 10.1038/modpathol.3880547

30 Sodir, N. M. et al. MYC Instructs and Maintains Pancreatic Adenocarcinoma Phenotype. Cancer Discovery 10, 588–607 (2020). 10.1158/2159-8290.CD-19-0435

31 Alison, M. R. The cellular origins of cancer with particular reference to the gastrointestinal tract. Int J Experimental Path 101, 132–151 (2020). 10.1111/iep.12364

32 Makohon-Moore, A. & Iacobuzio-Donahue, C. A. Pancreatic cancer biology and genetics from an evolutionary perspective. Nat Rev Cancer 16, 553–565 (2016). 10.1038/nrc.2016.66

33 Kopp, J. L. et al. Identification of Sox9-Dependent Acinar-to-Ductal Reprogramming as the Principal Mechanism for Initiation of Pancreatic Ductal Adenocarcinoma. Cancer Cell 22, 737–750 (2012). 10.1016/j.ccr.2012.10.025

34 Bailey, M. H. et al. Comprehensive Characterization of Cancer Driver Genes and Mutations. Cell 173, 371–385.e318 (2018). 10.1016/j.cell.2018.02.060

35 Malempati, S. et al. Aberrant stabilization of c-Myc protein in some lymphoblastic leukemias. Leukemia 20, 1572–1581 (2006). 10.1038/sj.leu.2404317

36 Escamilla-Powers, J. R. & Sears, R. C. A conserved pathway that controls c-Myc protein stability through opposing phosphorylation events occurs in yeast. J Biol Chem 282, 5432–5442 (2007). 10.1074/jbc.M611437200

37 Wang, X. et al. Phosphorylation Regulates c-Myc’s Oncogenic Activity in the Mammary Gland. Cancer Research 71, 925–936 (2011). 10.1158/0008-5472.CAN-10-1032

38 Zhang, X. et al. Mechanistic insight into Myc stabilization in breast cancer involving aberrant Axin1 expression. Proceedings of the National Academy of Sciences 109, 2790–2795 (2012). 10.1073/pnas.1100764108

39 Daniel, C. J. et al. Detection of Post-translational Modifications on MYC. Methods Mol Biol 2318, 69–85 (2021). 10.1007/978-1-0716-1476-1_5

40 Daniel, C. J., Zhang, X. & Sears, R. C. Detection of c-Myc protein-protein interactions and phosphorylation status by immunoprecipitation. Methods Mol Biol 1012, 65–76 (2013). 10.1007/978-1-62703-429-6_5

41 Murphy, D. J. et al. Distinct thresholds govern Myc’s biological output in vivo. Cancer Cell 14, 447–457 (2008). 10.1016/j.ccr.2008.10.018

42 Risom, T. et al. Deregulating MYC in a model of HER2+ breast cancer mimics human intertumoral heterogeneity. Journal of Clinical Investigation 130, 231–246 (2019). 10.1172/JCI126390

43 Doha, Z. O. et al. MYC Deregulation and PTEN Loss Model Tumor and Stromal Heterogeneity of Aggressive Triple-Negative Breast Cancer. Nat Commun 14, 5665 (2023). 10.1038/s41467-023-40841-6

44 Hingorani, S. R. et al. Trp53R172H and KrasG12D cooperate to promote chromosomal instability and widely metastatic pancreatic ductal adenocarcinoma in mice. Cancer Cell 7, 469–483 (2005). 10.1016/j.ccr.2005.04.023

45 Lee, A. Y. L. et al. Cell of origin affects tumour development and phenotype in pancreatic ductal adenocarcinoma. Gut 68, 487–498 (2019). 10.1136/gutjnl-2017-314426

46 Burlison, J. S., Long, Q., Fujitani, Y., Wright, C. V. & Magnuson, M. A. Pdx-1 and Ptf1a concurrently determine fate specification of pancreatic multipotent progenitor cells. Dev Biol 316, 74–86 (2008). 10.1016/j.ydbio.2008.01.011

47 Jennings, R. E. et al. Development of the human pancreas from foregut to endocrine commitment. Diabetes 62, 3514–3522 (2013). 10.2337/db12-1479

48 Dong, P. D., Provost, E., Leach, S. D. & Stainier, D. Y. Graded levels of Ptf1a differentially regulate endocrine and exocrine fates in the developing pancreas. Genes Dev 22, 1445–1450 (2008). 10.1101/gad.1663208

49 Hald, J. et al. Generation and characterization of Ptf1a antiserum and localization of Ptf1a in relation to Nkx6.1 and Pdx1 during the earliest stages of mouse pancreas development. J Histochem Cytochem 56, 587–595 (2008). 10.1369/jhc.2008.950675

50 Kawaguchi, Y. et al. The role of the transcriptional regulator Ptf1a in converting intestinal to pancreatic progenitors. Nat Genet 32, 128–134 (2002). 10.1038/ng959

51 Pan, F. C. et al. Spatiotemporal patterns of multipotentiality in Ptf1a-expressing cells during pancreas organogenesis and injury-induced facultative restoration. Development 140, 751–764 (2013). 10.1242/dev.090159

52 Kortlever, R. M. et al. Myc Cooperates with Ras by Programming Inflammation and Immune Suppression. Cell 171, 1301–1315.e1314 (2017). 10.1016/j.cell.2017.11.013

53 Tuveson, D. A. et al. Endogenous oncogenic K-ras(G12D) stimulates proliferation and widespread neoplastic and developmental defects. Cancer Cell 5, 375–387 (2004). 10.1016/s1535-6108(04)00085-6

54 Kiemen, A. L. et al. CODA: quantitative 3D reconstruction of large tissues at cellular resolution. Nat Methods 19, 1490–1499 (2022). 10.1038/s41592-022-01650-9

55 Bulle, A. & Lim, K.-H. Beyond just a tight fortress: contribution of stroma to epithelial-mesenchymal transition in pancreatic cancer. Sig Transduct Target Ther 5, 249 (2020). 10.1038/s41392-020-00341-1

56 Rebelo, R., Xavier, C. P. R., Giovannetti, E. & Vasconcelos, M. H. Fibroblasts in pancreatic cancer: molecular and clinical perspectives. Trends in Molecular Medicine 29, 439–453 (2023). 10.1016/j.molmed.2023.03.002

57 Ho, W. J., Jaffee, E. M. & Zheng, L. The tumour microenvironment in pancreatic cancer — clinical challenges and opportunities. Nat Rev Clin Oncol 17, 527–540 (2020). 10.1038/s41571-020-0363-5

58 Evan, G. I. et al. Re-engineering the Pancreas Tumor Microenvironment: A “Regenerative Program” Hacked. Clinical Cancer Research 23, 1647–1655 (2017). 10.1158/1078-0432.CCR-16-3275

59 Bardeesy, N. et al. *Smad4* is dispensable for normal pancreas development yet critical in progression and tumor biology of pancreas cancer. Genes Dev. 20, 3130–3146 (2006). 10.1101/gad.1478706

60 Cicenas, J. et al. KRAS, TP53, CDKN2A, SMAD4, BRCA1, and BRCA2 Mutations in Pancreatic Cancer. Cancers 9, 42 (2017). 10.3390/cancers9050042

61 Izeradjene, K. et al. KrasG12D and Smad4/Dpc4 Haploinsufficiency Cooperate to Induce Mucinous Cystic Neoplasms and Invasive Adenocarcinoma of the Pancreas. Cancer Cell 11, 229–243 (2007). 10.1016/j.ccr.2007.01.017

62 Oshima, M. et al. Immunohistochemically Detected Expression of 3 Major Genes (CDKN2A/p16, TP53, and SMAD4/DPC4) Strongly Predicts Survival in Patients With Resectable Pancreatic Cancer. Annals of Surgery 258, 336 (2013). 10.1097/SLA.0b013e3182827a65

63 Redston, M. S. et al. p53 mutations in pancreatic carcinoma and evidence of common involvement of homocopolymer tracts in DNA microdeletions. Cancer Research 54, 3025–3033 (1994).

64 Olive, K. P. et al. Mutant p53 Gain of Function in Two Mouse Models of Li-Fraumeni Syndrome. Cell 119, 847–860 (2004). 10.1016/j.cell.2004.11.004

65 Matsumoto, N. et al. Correlative Assessment of p53 Immunostaining Patterns and TP53 Mutation Status by Next-Generation Sequencing in High-Grade Endometrial Carcinomas. International Journal of Gynecological Pathology 42, 567 (2023). 10.1097/PGP.0000000000000930

66 Zhang, J. & Chen, X. Posttranscriptional Regulation of p53 and its Targets by RNABinding Proteins. Current Molecular Medicine 8, 845–849 10.2174/156652408786733748

67 Al-Khalaf, H. H. & Aboussekhra, A. p16 Controls p53 Protein Expression Through miR- dependent Destabilization of MDM2. Molecular Cancer Research 16, 1299–1308 (2018). 10.1158/1541-7786.MCR-18-0017

68 Niknafs, N. et al. Characterization of genetic subclonal evolution in pancreatic cancer mouse models. Nat Commun 10, 5435 (2019). 10.1038/s41467-019-13100-w

69 Harada, T. et al. Genome-wide DNA copy number analysis in pancreatic cancer using high- density single nucleotide polymorphism arrays. Oncogene 27, 1951–1960 (2008). 10.1038/sj.onc.1210832

70 Bailey, P. et al. Genomic analyses identify molecular subtypes of pancreatic cancer. Nature 531, 47–52 (2016). 10.1038/nature16965

71 Gao, F.-Y., Li, X.-T., Xu, K., Wang, R.-T. & Guan, X. -x. c-MYC mediates the crosstalk between breast cancer cells and tumor microenvironment. Cell Commun Signal 21, 28 (2023). 10.1186/s12964-023-01043-1

72 Thege, F. I. et al. A Programmable *In Vivo* CRISPR Activation Model Elucidates the Oncogenic and Immunosuppressive Functions of MYC in Lung Adenocarcinoma. Cancer Research 82, 2761–2776 (2022). 10.1158/0008-5472.CAN-21-4009

73 Blise, K. E., Sivagnanam, S., Banik, G. L., Coussens, L. M. & Goecks, J. Single-cell spatial architectures associated with clinical outcome in head and neck squamous cell carcinoma. NPJ Precis Oncol 6, 10 (2022). 10.1038/s41698-022-00253-z

74 Jackson, H. W. et al. The single-cell pathology landscape of breast cancer. Nature 578, 615–620 (2020). 10.1038/s41586-019-1876-x

75 Hänzelmann, S., Castelo, R. & Guinney, J. GSVA: gene set variation analysis for microarray and RNA-Seq data. BMC Bioinformatics 14, 7 (2013). 10.1186/1471-2105-14-7

76 Liberzon, A. et al. The Molecular Signatures Database (MSigDB) hallmark gene set collection. Cell Syst 1, 417–425 (2015). 10.1016/j.cels.2015.12.004

77 Rashid, N. U. et al. Purity Independent Subtyping of Tumors (PurIST), A Clinically Robust, Single-sample Classifier for Tumor Subtyping in Pancreatic Cancer. Clin Cancer Res 26, 82–92 (2020). 10.1158/1078-0432.CCR-19-1467

78 Li, Y. et al. Patterns of somatic structural variation in human cancer genomes. Nature 578, 112–121 (2020). 10.1038/s41586-019-1913-9

79 Subramanian, A. et al. Gene set enrichment analysis: A knowledge-based approach for interpreting genome-wide expression profiles. Proceedings of the National Academy of Sciences 102, 15545–15550 (2005). 10.1073/pnas.0506580102

80 Mi, H., Muruganujan, A., Ebert, D., Huang, X. & Thomas, P. D. PANTHER version 14: more genomes, a new PANTHER GO-slim and improvements in enrichment analysis tools. Nucleic Acids Research 47, D419–D426 (2019). 10.1093/nar/gky1038

81 Thomas, P. D. et al. PANTHER: Making genome-scale phylogenetics accessible to all. Protein Science 31, 8–22 (2022). 10.1002/pro.4218

82 Durinck, S., Spellman, P. T., Birney, E. & Huber, W. Mapping identifiers for the integration of genomic datasets with the R/Bioconductor package biomaRt. Nat Protoc 4, 1184–1191 (2009). 10.1038/nprot.2009.97

83 Dyer, S. C. et al. Ensembl 2025. Nucleic Acids Res 53, D948–D957 (2025). 10.1093/nar/gkae1071

84 Durinck, S. et al. BioMart and Bioconductor: a powerful link between biological databases and microarray data analysis. Bioinformatics 21, 3439–3440 (2005). 10.1093/bioinformatics/bti525

85 Zhang, Y., Parmigiani, G. & Johnson, W. E. ComBat-seq: batch effect adjustment for RNA-seq count data. NAR Genom Bioinform 2, lqaa078 (2020). 10.1093/nargab/lqaa078

86 Moffitt, R. A. et al. Virtual microdissection identifies distinct tumor- and stroma-specific subtypes of pancreatic ductal adenocarcinoma. Nat Genet 47, 1168–1178 (2015). 10.1038/ng.3398

87 Collisson, E. A., Bailey, P., Chang, D. K. & Biankin, A. V. Molecular subtypes of pancreatic cancer. Nat Rev Gastroenterol Hepatol 16, 207–220 (2019). 10.1038/s41575-019-0109-y

88 Pishvaian, M. J. & Brody, J. R. Therapeutic Implications of Molecular Subtyping for Pancreatic Cancer. Oncology (Williston Park) 31, 159–166, 168 (2017).

89 Cancer of the Pancreas - Cancer Stat Facts. SEER

90 Stoop, T. F. et al. Pancreatic cancer. Lancet 405, 1182–1202 (2025). 10.1016/S0140-6736(25)00261-2

91 Aguirre, A. J. et al. Activated Kras and *Ink4a/Arf* deficiency cooperate to produce metastatic pancreatic ductal adenocarcinoma. Genes Dev. 17, 3112–3126 (2003). 10.1101/gad.1158703

92 Herreros-Villanueva, M., Hijona, E., Cosme, A. & Bujanda, L. Mouse models of pancreatic cancer. World J Gastroenterol 18, 1286–1294 (2012). 10.3748/wjg.v18.i12.1286

93 Hingorani, S. R. et al. Preinvasive and invasive ductal pancreatic cancer and its early detection in the mouse. Cancer Cell 4, 437–450 (2003). 10.1016/S1535-6108(03)00309-X

94 Skoulidis, F. et al. Germline Brca2 heterozygosity promotes Kras(G12D) -driven carcinogenesis in a murine model of familial pancreatic cancer. Cancer Cell 18, 499–509 (2010). 10.1016/j.ccr.2010.10.015

95 Fukushima, N. et al. Aberrant methylation of preproenkephalin and p16 genes in pancreatic intraepithelial neoplasia and pancreatic ductal adenocarcinoma. Am J Pathol 160, 1573–1581 (2002). 10.1016/S0002-9440(10)61104-2

96 Wilentz, R. E. et al. Loss of expression of Dpc4 in pancreatic intraepithelial neoplasia: evidence that DPC4 inactivation occurs late in neoplastic progression. Cancer Res 60, 2002–2006 (2000).

97 Yang, K., Li, X. & Xie, K. Senescence program and its reprogramming in pancreatic premalignancy. Cell Death Dis 14, 1–11 (2023). 10.1038/s41419-023-06040-3

98 Davalos, A. R., Coppe, J.-P., Campisi, J. & Desprez, P.-Y. Senescent cells as a source of inflammatory factors for tumor progression. Cancer Metastasis Rev 29, 273–283 (2010). 10.1007/s10555-010-9220-9

99 Moir, J. A. G., White, S. A. & Mann, J. Arrested development and the great escape--the role of cellular senescence in pancreatic cancer. Int J Biochem Cell Biol 57, 142–148 (2014). 10.1016/j.biocel.2014.10.018

100 Balachandran, V. P., Beatty, G. L. & Dougan, S. K. Broadening the Impact of Immunotherapy to Pancreatic Cancer: Challenges and Opportunities. Gastroenterology 156, 2056–2072 (2019). 10.1053/j.gastro.2018.12.038

101 Mi, H. et al. Quantitative Spatial Profiling of Immune Populations in Pancreatic Ductal Adenocarcinoma Reveals Tumor Microenvironment Heterogeneity and Prognostic Biomarkers. Cancer Research 82, 4359–4372 (2022). 10.1158/0008-5472.CAN-22-1190

102 Evans, R. A. et al. Lack of immunoediting in murine pancreatic cancer reversed with neoantigen. JCI Insight 1 (2016). 10.1172/jci.insight.88328

103 Barilla, R. M. et al. Specialized dendritic cells induce tumor-promoting IL-10+IL-17+ FoxP3neg regulatory CD4+ T cells in pancreatic carcinoma. Nat Commun 10, 1424 (2019). 10.1038/s41467-019-09416-2

104 Hegde, S. et al. Dendritic Cell Paucity Leads to Dysfunctional Immune Surveillance in Pancreatic Cancer. Cancer Cell 37, 289–307.e289 (2020). 10.1016/j.ccell.2020.02.008

105 Lomberk, G. et al. Distinct epigenetic landscapes underlie the pathobiology of pancreatic cancer subtypes. Nat Commun 9, 1978 (2018). 10.1038/s41467-018-04383-6

106 Cortes-Ciriano, I. et al. Comprehensive analysis of chromothripsis in 2,658 human cancers using whole-genome sequencing. Nat Genet 52, 331–341 (2020). 10.1038/s41588-019-0576-7

107 Collisson, E. A. et al. Subtypes of pancreatic ductal adenocarcinoma and their differing responses to therapy. Nat Med 17, 500–503 (2011). 10.1038/nm.2344

108 Moffitt, R. A. et al. Virtual microdissection identifies distinct tumor- and stroma-specific subtypes of pancreatic ductal adenocarcinoma. Nat Genet 47, 1168–1178 (2015). 10.1038/ng.3398

109 Chan-Seng-Yue, M. et al. Transcription phenotypes of pancreatic cancer are driven by genomic events during tumor evolution. Nat Genet 52, 231–240 (2020). 10.1038/s41588-019-0566-9

110 He, M., Henderson, M., Muth, S., Murphy, A. & Zheng, L. Preclinical mouse models for immunotherapeutic and non-immunotherapeutic drug development for pancreatic ductal adenocarcinoma. Ann Pancreat Cancer 3 (2020). 10.21037/apc.2020.03.03

111 Kawaguchi, Y. et al. The role of the transcriptional regulator Ptf1a in converting intestinal to pancreatic progenitors. Nat Genet 32, 128–134 (2002). 10.1038/ng959

112 Greenwald, N. F. et al. Whole-cell segmentation of tissue images with human-level performance using large-scale data annotation and deep learning. Nat Biotechnol 40, 555–565 (2022). 10.1038/s41587-021-01094-0

113 Tsujikawa, T. et al. Quantitative Multiplex Immunohistochemistry Reveals Myeloid-Inflamed Tumor-Immune Complexity Associated with Poor Prognosis. Cell Reports 19, 203–217 (2017). 10.1016/j.celrep.2017.03.037

114 Liudahl, S. M. et al. Leukocyte Heterogeneity in Pancreatic Ductal Adenocarcinoma: Phenotypic and Spatial Features Associated with Clinical Outcome. Cancer Discovery 11, 2014–2031 (2021). 10.1158/2159-8290.CD-20-0841

115 Banik, G. et al. High-dimensional multiplexed immunohistochemical characterization of immune contexture in human cancers. Methods Enzymol 635, 1–20 (2020). 10.1016/bs.mie.2019.05.039

116 Means, C. et al. Tumor immune microenvironment characteristics of papillary thyroid carcinoma are associated with histopathological aggressiveness and BRAF mutation status. Head & Neck 41, 2636–2646 (2019). 10.1002/hed.25740

117 Reinhard, E., Adhikhmin, M., Gooch, B. & Shirley, P. Color transfer between images. IEEE Comput. Grap. Appl. 21, 34–41 (2001). 10.1109/38.946629

118. Schindelin, J. et al. Fiji: an open-source platform for biological-image analysis. Nat Methods 9, 676-682 (2012). 10.1038/nmeth.2019

119 Carpenter, A. E. et al. CellProfiler: image analysis software for identifying and quantifying cell phenotypes. Genome Biol 7, R100 (2006). 10.1186/gb-2006-7-10-r100

120 Gao, J. et al. Integrative Analysis of Complex Cancer Genomics and Clinical Profiles Using the cBioPortal. Sci. Signal. 6 (2013). 10.1126/scisignal.2004088

121 Cerami, E. et al. The cBio Cancer Genomics Portal: An Open Platform for Exploring Multidimensional Cancer Genomics Data. Cancer Discovery 2, 401–404 (2012). 10.1158/2159-8290.CD-12-0095

122 Li, H. & Durbin, R. Fast and accurate short read alignment with Burrows–Wheeler transform. Bioinformatics 25, 1754–1760 (2009). 10.1093/bioinformatics/btp324

123 Langmead, B., Trapnell, C., Pop, M. & Salzberg, S. L. Ultrafast and memory-efficient alignment of short DNA sequences to the human genome. Genome Biol 10, R25 (2009). 10.1186/gb-2009-10-3-r25

124 Adalsteinsson, V. A. et al. Scalable whole-exome sequencing of cell-free DNA reveals high concordance with metastatic tumors. Nat Commun 8, 1324 (2017). 10.1038/s41467-017-00965-y

125 Garcia, M. et al. Sarek: A portable workflow for whole-genome sequencing analysis of germline and somatic variants. F1000Res 9, 63 (2020). 10.12688/f1000research.16665.2

126 Ewels, P. A. et al. The nf-core framework for community-curated bioinformatics pipelines. Nat Biotechnol 38, 276–278 (2020). 10.1038/s41587-020-0439-x

127 Wardell, C. P., Ashby, C. & Bauer, M. A. FiNGS: high quality somatic mutations using filters for next generation sequencing. BMC Bioinformatics 22, 77 (2021). 10.1186/s12859-021-03995-y

128 Bray, N. L., Pimentel, H., Melsted, P. & Pachter, L. Near-optimal probabilistic RNA-seq quantification. Nat Biotechnol 34, 525–527 (2016). 10.1038/nbt.3519

129 Köster, J. & Rahmann, S. Snakemake—a scalable bioinformatics workflow engine. Bioinformatics 28, 2520–2522 (2012). 10.1093/bioinformatics/bts480

130 R: A Language and Environment for Statistical Computing (R Foundation for Statistical Computing, Vienna, Austria, 2021).

131 Love, M. I., Huber, W. & Anders, S. Moderated estimation of fold change and dispersion for RNA-seq data with DESeq2. Genome Biol 15, 550 (2014). 10.1186/s13059-014-0550-8

132 Soneson, C., Love, M. I. & Robinson, M. D. Differential analyses for RNA-seq: transcript-level estimates improve gene-level inferences. F1000Res 4, 1521 (2016). 10.12688/f1000research.7563.2

133 Wickham, H. ggplot2: Elegant Graphics for Data Analysis., (Springer-Verlag, 2016).

134 pheatmap: Pretty Heatmaps (2019).

135 Ashburner, M. et al. Gene Ontology: tool for the unification of biology. Nat Genet 25, 25–29 (2000). 10.1038/75556

136 The Gene Ontology, C. et al. The Gene Ontology resource: enriching a GOld mine. Nucleic Acids Research 49, D325–D334 (2021). 10.1093/nar/gkaa1113

137 Mi, H. et al. Protocol Update for large-scale genome and gene function analysis with the PANTHER classification system (v.14.0). *Nat Protoc* 14, 703-721 (2019). 10.1038/s41596-019-0128-8

138 Dolgalev, I. msigdbr: MSigDB Gene Sets for Multiple Organisms in a Tidy Data Format. (2025).

139 Mootha, V. K. et al. PGC-1α-responsive genes involved in oxidative phosphorylation are coordinately downregulated in human diabetes. Nat Genet 34, 267–273 (2003). 10.1038/ng1180

140 Liberzon, A. et al. The Molecular Signatures Database Hallmark Gene Set Collection. Cell Systems 1, 417–425 (2015). 10.1016/j.cels.2015.12.004

141 Therneau, T. A package for survival analysis in R. (2024).

142 Alboukadel Kassambara, M. K., Przemyslaw Biecek. survminer: Drawing Survival Curves using ’ggplot2’. (2024).

143 Boj, Sylvia F. et al. Organoid Models of Human and Mouse Ductal Pancreatic Cancer. Cell 160, 324–338 (2015). 10.1016/j.cell.2014.12.021

144 Aiello, N. M., Rhim, A. D. & Stanger, B. Z. Orthotopic Injection of Pancreatic Cancer Cells. Cold Spring Harb Protoc 2016, pdb.prot078360 (2016). 10.1101/pdb.prot078360

145 Borsoi, C. et al. Gemcitabine enhances the transport of nanovector-albumin-bound paclitaxel in gemcitabine-resistant pancreatic ductal adenocarcinoma. Cancer Letters 403, 296–304 (2017). 10.1016/j.canlet.2017.06.026

146 Horiuchi, T. et al. New treatment strategy with nuclear factor-κB inhibitor for pancreatic cancer. Journal of Surgical Research 206, 1-8 (2016). 10.1016/j.jss.2016.06.047

147 Sasaki, T. et al. Targeting claudin-4 enhances chemosensitivity of pancreatic ductal carcinomas. Cancer Medicine 8, 6700–6708 (2019). 10.1002/cam4.2547

