## supplementary figures and tables for "Myc and Kras cooperate in adult acinar cells to drive phenotypic heterogeneity, metastasis, and therapeutic resistance in a novel pancreatic cancer mouse model"

### Supplementary Figure 1

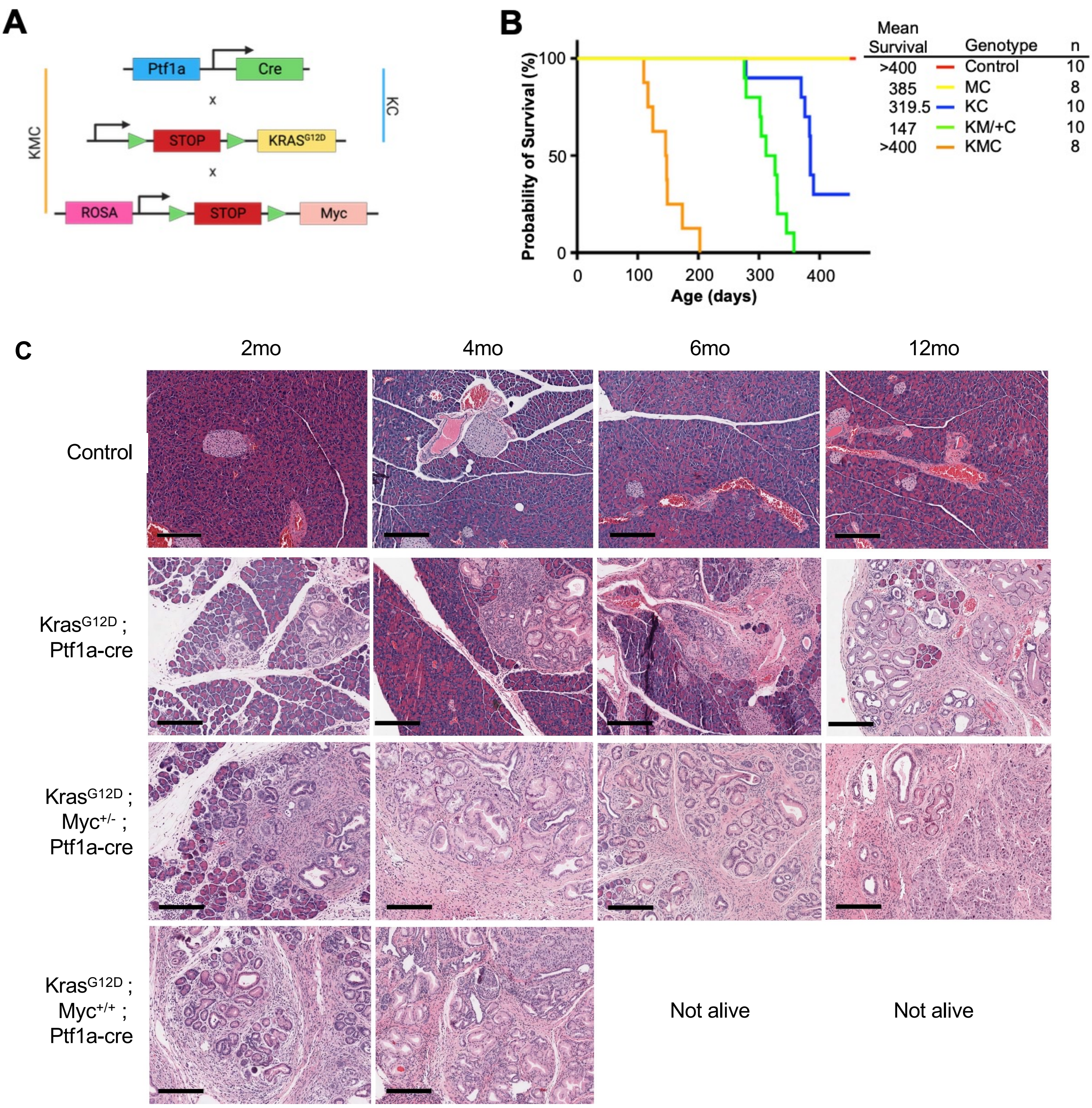

### Supplementary Figure 2

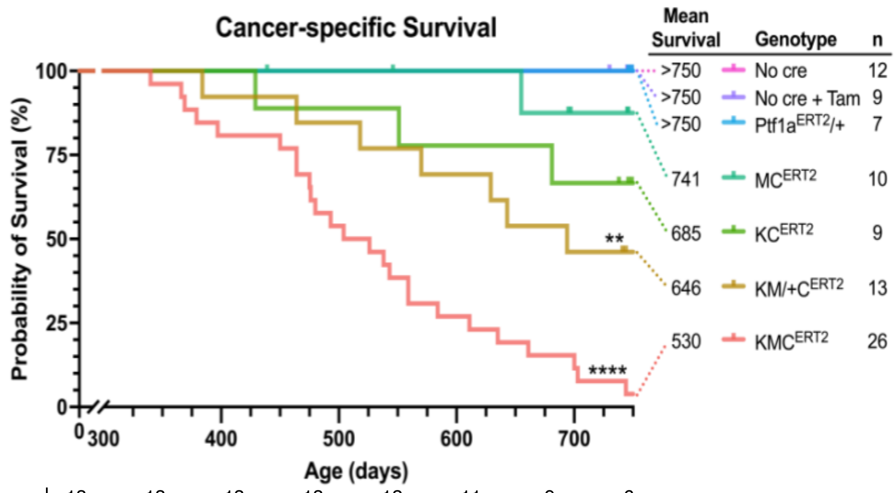

| Genotype | 0 | 100 | 200 | 300 | 400 | 500 | 600 | 700 |
|---|---|---|---|---|---|---|---|---|
| | 13 | 13 | 13 | 13 | 12 | 11 | 9 | 6 |
| | 9 | 9 | 9 | 9 | 9 | 8 | 7 | 6 |
| | 10 | 10 | 10 | 10 | 10 | 9 | 8 | 5 |
| | 7 | 7 | 7 | 7 | 7 | 7 | 7 | 7 |
| | 9 | 9 | 9 | 9 | 9 | 9 | 9 | 9 |
| | 12 | 12 | 12 | 12 | 12 | 12 | 12 | 12 |

Time (days)

529.6
646.3
684.5
740.5
> 750
> 750
> 750

### (Supplementary) Table 1

| Autochthonous Tumor Endpoint Data | | | | | | | Omics Data Availability | | | Derived Models | |
|---|---|---|---|---|---|---|---|---|---|---|---|
| Mouse ID | Sex | Genotype | Tumor subtype | Overall survival (days) | Relative pancreas weight (% bodyweight) | Macro metastases noted on necropsy | RNAseq | WGS | WES | Cell line grown | Orthotopic growth |
| 011T | M | $KMC^{ERT2}$ | Not assessed | 635 | 7.36% | - | - | - | - | + | Not assessed |
| 078J | F | $KMC^{ERT2}$ | Squamous | 700 | Not available | + | + | + | - | - | N/A |
| 106L | F | $KMC^{ERT2}$ | Not assessed | 584 | 6.90% | - | - | - | - | + | Not assessed |
| 136J | F | $KMC^{ERT2}$ | Squamous | 369 | 9.35% | + | + | + | - | - | N/A |
| 167J | F | $KMC^{ERT2}$ | Squamous | 493 | 5.82% | + | + | + | - | - | N/A |
| 171J | M | $KMC^{ERT2}$ | Squamous | 611 | 5.98% | + | + | + | + | + | Consistent |
| 222J | M | $KM/+C^{ERT2}$ | Pancreatic Progenitor | 629 | 2.80% | - | + | + | - | - | N/A |
| 255J | F | $KMC^{ERT2}$ | Squamous | 480 | 8.01% | + | + | + | + | + | Inconsistent |
| 301J | M | $KMC^{ERT2}$ | Pancreatic Progenitor | 559 | 5.34% | - | + | + | + | + | Consistent |
| 303J | M | $KMC^{ERT2}$ | Pancreatic Progenitor | 366 | 6.73% | + | + | + | + | + | Consistent |
| 304J | F | $KM/+C^{ERT2}$ | Pancreatic Progenitor | 464 | 7.11% | - | + | + | - | - | N/A |
| 313J | F | $KMC^{ERT2}$ | Pancreatic Progenitor | 379 | 6.76% | - | + | + | - | - | N/A |
| 314J | M | $KMC^{ERT2}$ | Squamous | 504 | 8.10% | + | + | + | + | + | Consistent |
| 352J | M | $KM/+C^{ERT2}$ | Pancreatic Progenitor | 518 | 5.04% | - | + | + | - | - | N/A |
| 369J | F | $KM/+C^{ERT2}$ | Pancreatic Progenitor | 384 | 5.70% | - | + | + | - | - | N/A |
| 808J | F | $KM/+C^{ERT2}$ | Not assessed | 543 | 5.64% | - | - | - | - | + | Not assessed |

### Supplementary Figure 3

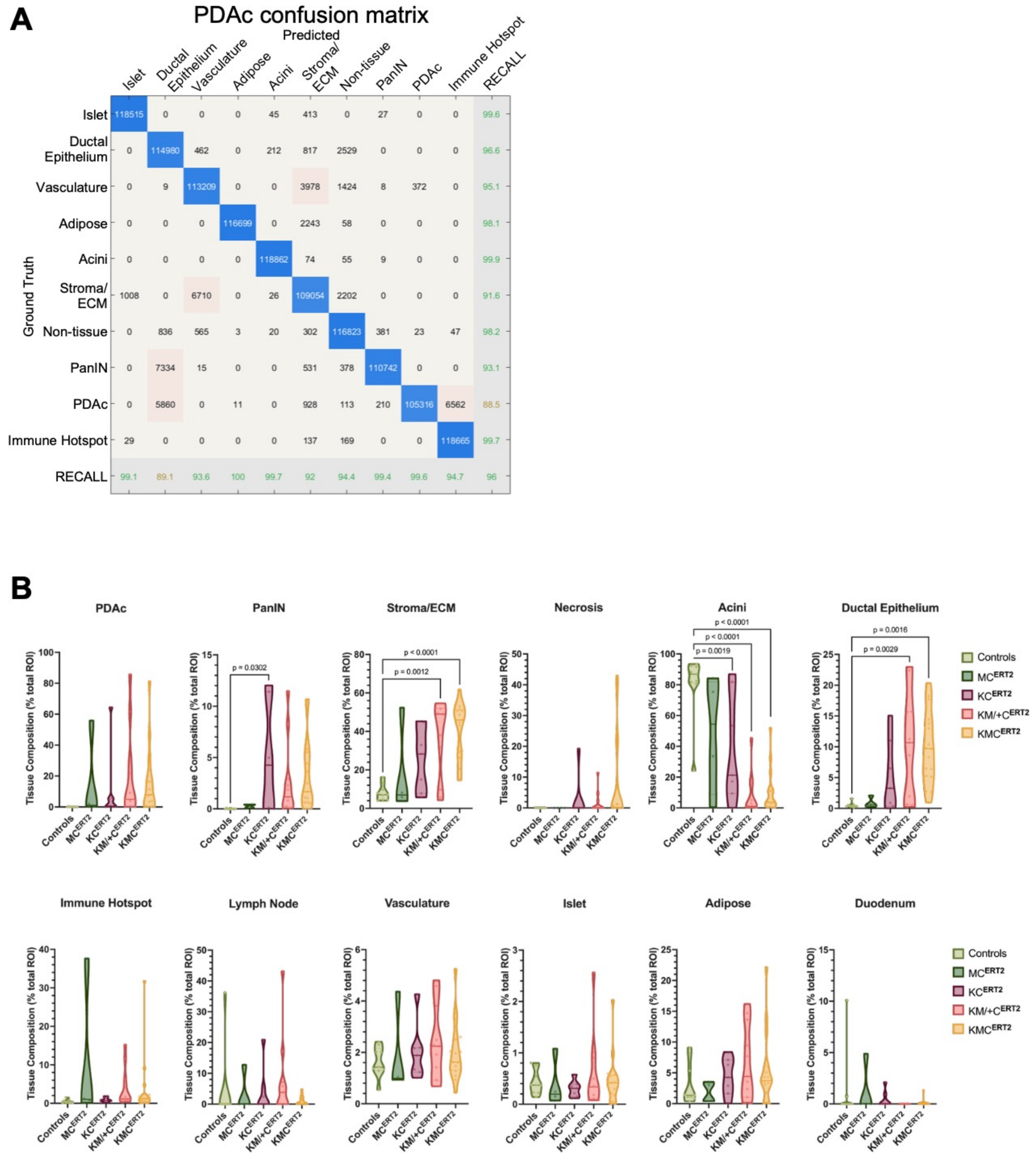

Supplementary Figure 3. CODA pancreatic tissue composition analysis

### Supplementary Table 2

| Sample ID | High Probable Impact (# [name]) | Moderate Probable Impact (#) | Total non-syn SNVs (#) |
|---|---|---|---|
| 171J | 1 (FGFR2) | 19 | 27 |
| 301J | 1 (ITGA2) | 32 | 38 |
| 303J | 0 | 24 | 31 |
| 314J | 3 (FRAS1, MYO15, TLE6) | 23 | 37 |

### Supplementary Figure 4

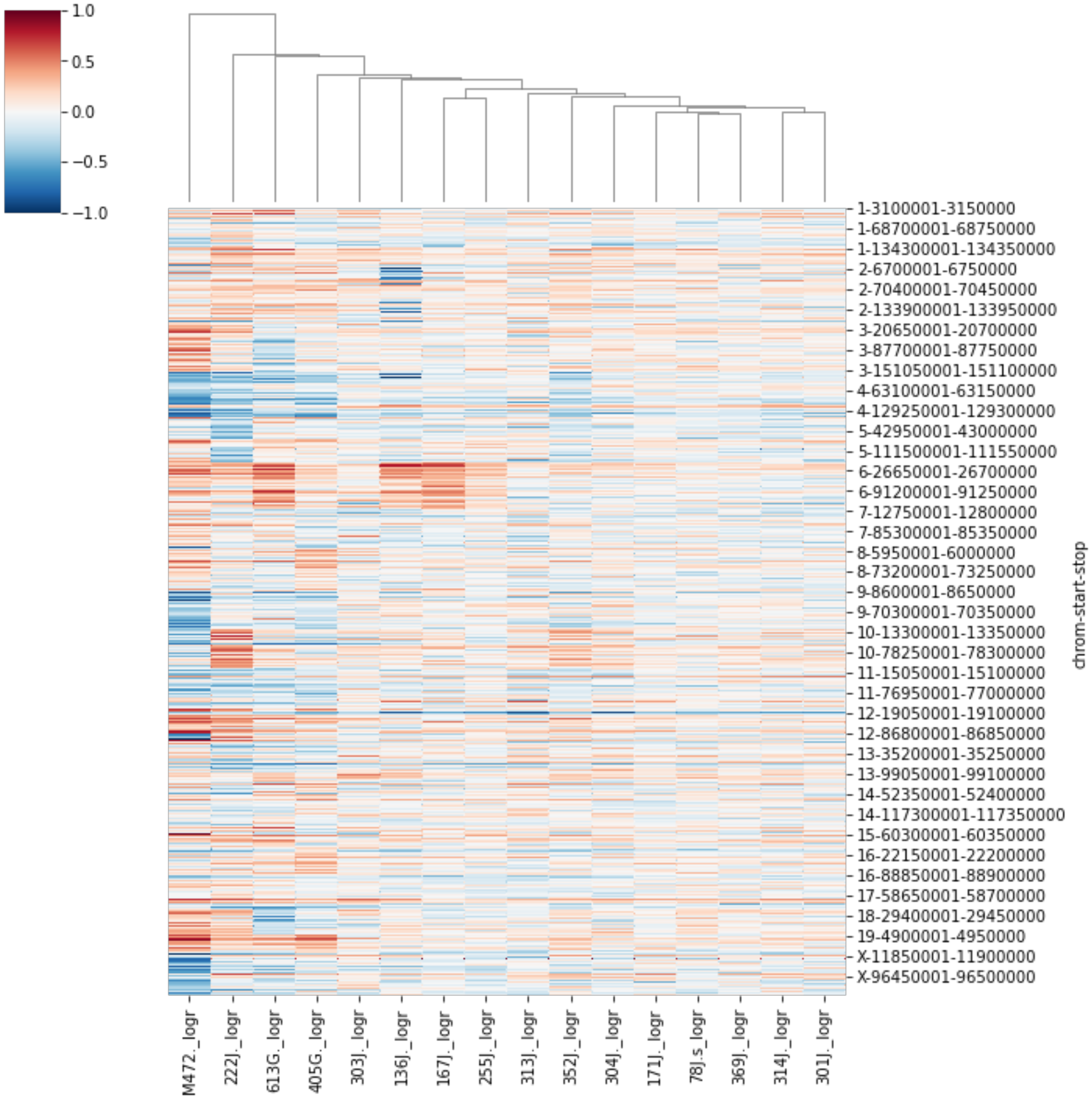

### Supplementary Table 3

| | Cycle 1 (Pre-Antigen Retrieval) | | Cycle 2 | Cycle 3 | Cycle 4 | Cycle 5 | Cycle 6 | Cycle 7 |
|---|---|---|---|---|---|---|---|---|
| | Round 1 | Round 1 | Round 1 | Round 1 | Round 1 | Round 1 | Round 1 | Round 1 |
| **Primary Antibody or Stain** | **CSF-1R** | **Hematoxylin** | **CD11c** | **CD4** | **BTK** | **PDL1** | **CD3** | **CD206** |
| **Clone** | E2412 | N/A | D1V9Y | D7D2Z | Polyclonal | E1L3N | SP7 | Polyclonal |
| **Vendor** | Santa Cruz | Dako | Cell Signaling | Cell Signaling | LS Bio | Cell Signaling | Thermo | Abcam |
| **Catalog #** | sc-692 | S3301 | 97585S | 25229 | LS-C180161 | 13684S | RM-9107-S | 64693 |
| **Host Species** | Rabbit | N/A | Rabbit | Rabbit | Rabbit | Rabbit | Rabbit | Rabbit |
| **Dilution** | 1:500 | N/A | 1:100 | 1:100 | 1:200 | 1:50 | 1:300 | 1:1000 |
| **Incubation Time** | 1 hr @ RT | 1 min @ RT | 1 hr @ RT | 4 hr @ RT | 1 hr @ RT | O/N @ 4°C | 1 hr @ RT | O/N @ 4°C |
| **AEC Development Time** | 40 min | N/A | 25 min | 40 min | 13 min | 25 min | 27 min | 15 min |
| | Round 2 | | | Round 2 | Round 2 | Round 2 | Round 2 | Round 2 |
| **Primary Antibody or Stain** | **F4/80** | | | **MHC II** | **CD45** | **CD8** | **CD207** | **B220** |
| **Clone** | Cl:A3-1 | | | M5/114.15.2 | 30-F11 | 4SM15 | eBioRMUL.2 | R13-6B2 |
| **Vendor** | BioRad | | | eBioscience | eBioscience | eBiosicence | eBioscience | BD |
| **Catalog #** | MCA497G | | | eBi14-5321 | 14-0451-82 | 14-0808-82 | 14-2073-82 | 550286 |
| **Host Species** | Rat | | | Rat | Rat | Rat | Rat | Rat |
| **Dilution** | 1:200 | | | 1:100 | 1:50 | 1: 100 | 1:100 | 1:100 |
| **Incubation Time** | 1 hr @ RT | | | 4 hr @ RT | 1 hr @ RT | O/N @ 4°C | 1 hr @ RT | O/N @ 4°C |
| **AEC Development Time** | 40 min | | | 14.5 min | 15 min | 25 min | 19 min | 40 min |

| | Cycle 8 | Cycle 9 | Cycle 10 | Cycle 11 | Cycle 12 | Cycle 13 | Cycle 14 | Cycle 15 |
|---|---|---|---|---|---|---|---|---|
| | Round 1 | Round 1 | Round 1 | Round 1 | Round 1 | Round 1 | Round 1 | Round 1 |
| **Primary Antibody or Stain** | **RORgt** | **GATA3** | **CD11b** | **TCF1/TCF7** | **TIM3** | **EOMES (TBR2)** | **Granzyme B** | **Ki67** |
| **Clone** | EPR20006 | EPR16651 | EPR1334 | C63D9 | D3M9R | EPR19012 | Polyclonal | Polyclonal |
| **Vendor** | Abcam | Abcam | Abcam | Cell Signaling | Cell Signalling | Abcam | Abcam | Abcam |
| **Catalog #** | ab207082 | 199428 | 133357 | 2203S | 83882S | 183991 | 4059 | 15580 |
| **Host Species** | Rabbit | Rabbit | Rabbit | Rabbit | Rabbit | Rabbit | Rabbit | Rabbit |
| **Dilution** | 1:100 | 1:100 | 1:3000 | 1:100 | 1:200 | 1:1000 | 1:200 | 1:5000 |
| **Incubation Time** | 1 hr @ RT | 30 min @ RT | 1 hr @ RT | 30 min @ RT | 30 min @ RT | 1 hr @ RT | O/N @ 4°C | 30 min @ RT |
| **AEC Development Time** | 20 min | 25 min | 10 min | 15 min | 25 min | 25 min | 10 min | 17 min |
| | Round 2 | | | | | | Round 2 | Round 2 |
| **Primary Antibody or Stain** | **Foxp3** | | | | | | **Ly6G** | **Pan Cytokeratin** |
| **Clone** | FJK16S | | | | | | 1A8 | AE1/1E3 |
| **Vendor** | eBioscience | | | | | | eBioscience | Abcam |
| **Catalog #** | 14-5773-82 | | | | | | 551459 | ab27988 |
| **Host Species** | Rat | | | | | | Rat | MOUSE |
| **Dilution** | 1:100 | | | | | | 1:200 | 1:100 |
| **Incubation Time** | 30 min @ RT | | | | | | O/N @ 4°C | 30 min @ RT |
| **AEC Development Time** | 30 min | | | | | | 20 min | 10 min |

### Supplementary Table 4

| | |
|---|---|
| **Immune cell type identification by marker expression** | |
| Lineage | Identification (All populations are $CD45^{+}$) |
| Th0 (naïve) helper T cells | $CD3^{+}$ $CD4^{+}$ $CD8^{-}$ $Foxp3^{-}$ $RORgt^{-}$ $Tbet^{-}$ $GATA3^{-}$ |
| Regulatory T cells (Treg) | $CD3^{+}$ $CD4^{+}$ $CD8^{-}$ $Foxp3^{+}$ |
| Th17 helper T cells | $CD3^{+}$ $CD4^{+}$ $CD8^{-}$ $RORgt^{+}$ |
| Th2 helper T cells | $CD3^{+}$ $CD4^{+}$ $CD8^{-}$ $GATA3^{+}$ |
| $CD8^{+}$ T lymphocytes (all) | $CD3^{+}$ $CD8^{+}$ |
| $CD8^{+}$ T lymphocytes (memory) | $CD3^{+}$ $CD8^{+}$ $EOMES^{+}$ |
| B cells | $CD3^{-}$ B220+ |
| Granulocytes | $CD3/B220^{-}$ $Ly6G^{+}$ |
| Pro-tumor TAM | $CD3/B220/Ly6G^{-}$ $F4/80^{+}$ $CSF1R^{+}$ $CD206^{+}$ |
| Re-programmed TAM | $CD3/B220/Ly6G^{-}$ $F4/80^{+}$ $CSF1R^{+}$ $CD206^{-}$ $MHCII^{+}$ |
| Other pro-tumor myeloid | $CD3/B220/Ly6G^{-}$ $F4/80^{+}$ $CSF1R^{-}$ $CD206^{+}$ |
| Other myeloid | $CD3/B220/Ly6G^{-}$ $F4/80^{+}$ $CSF1R^{-}$ $CD206^{-}$ |
| Conventional DC (DC1) | $CD3/B220/Ly6G^{-}$ $F4/80^{-}$ $CD11c^{+}$ $MHCII^{+}$ $CD11b^{+}$ |
| Cross presenting DC (DC2) | $CD3/B220/Ly6G^{-}$ $F4/80^{-}$ $CD11c^{+}$ $MHCII^{+}$ $CD11b^{-}$ CD207+ |
| **Interrogation of functional state of immune cells** | |
| Marker | Classification |
| Proliferation | Ki67 |
| Cytotoxicity | Granzyme-B+ |
| T cell activation | TCF1/TCF7 |
| Peripheral tolerance | PD-L1 |
| **Identification of non-immune cells** | |
| Cell type | Identification (All populations are $CD45^{-}$) |
| Activated fibroblast + SMCs | $\alpha SMA^{+}$ |
| Endothelial cells | $CD31^{+}$ |

### Supplementary Figure 5

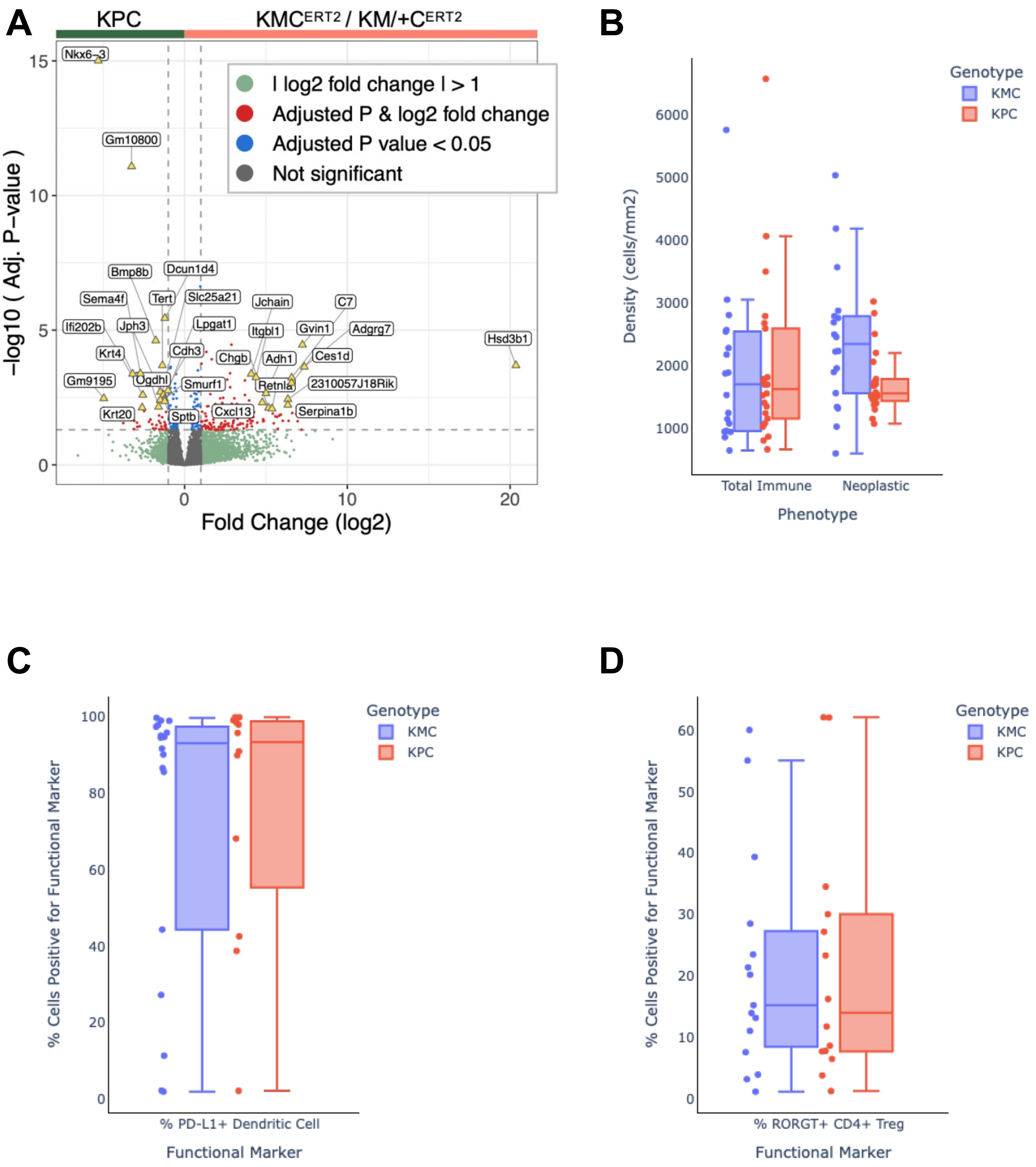

**Supplementary Figure 6.** KMC$^{ERT2}$ PDAc model displays intertumoral transcriptional and phenotypic heterogeneity.

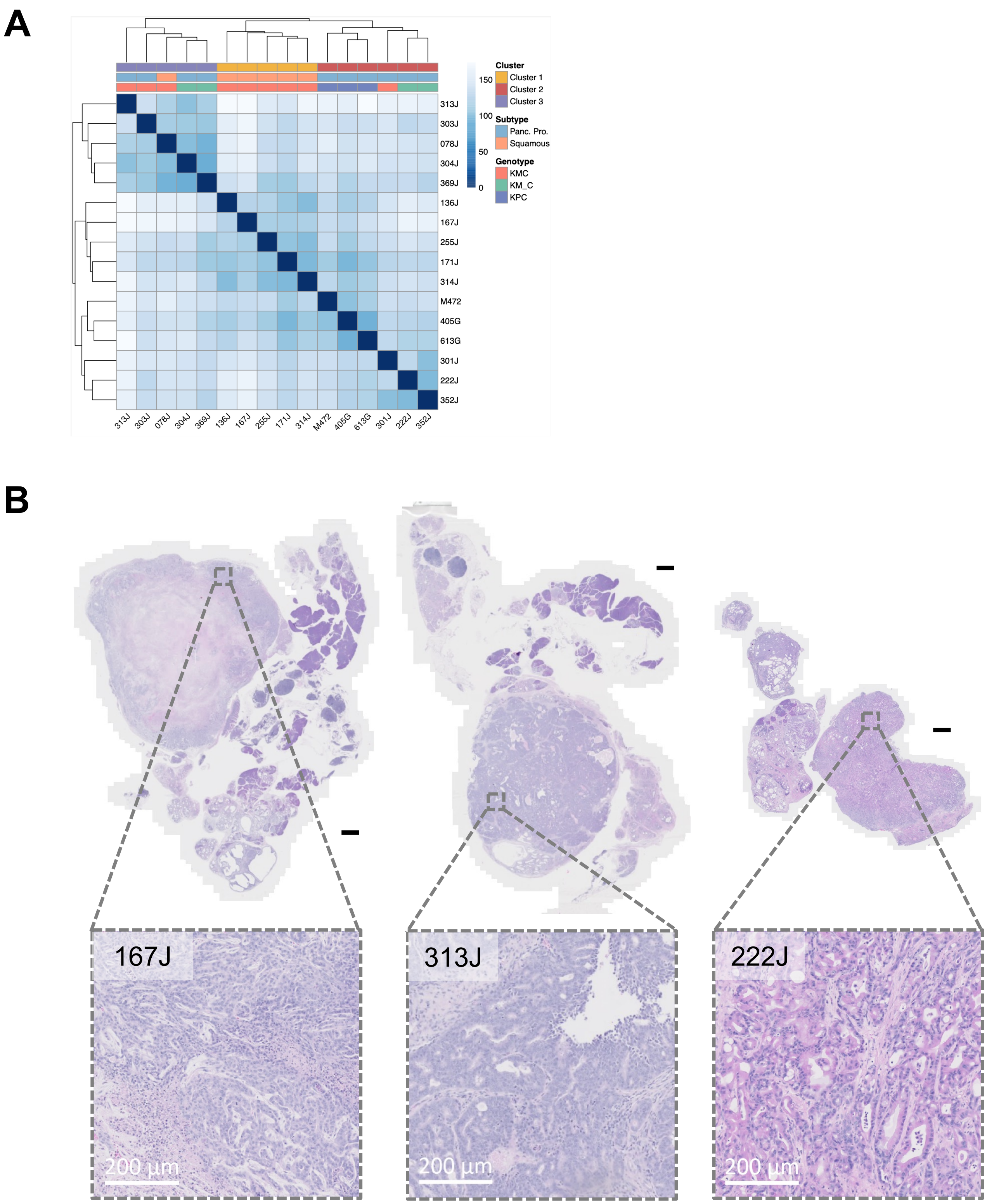

**Supplementary Figure 7.** Cluster 1 tumors are upregulated in cell cycle, DNA replication and repair biological process GO Terms and GSEA hallmark pathways.

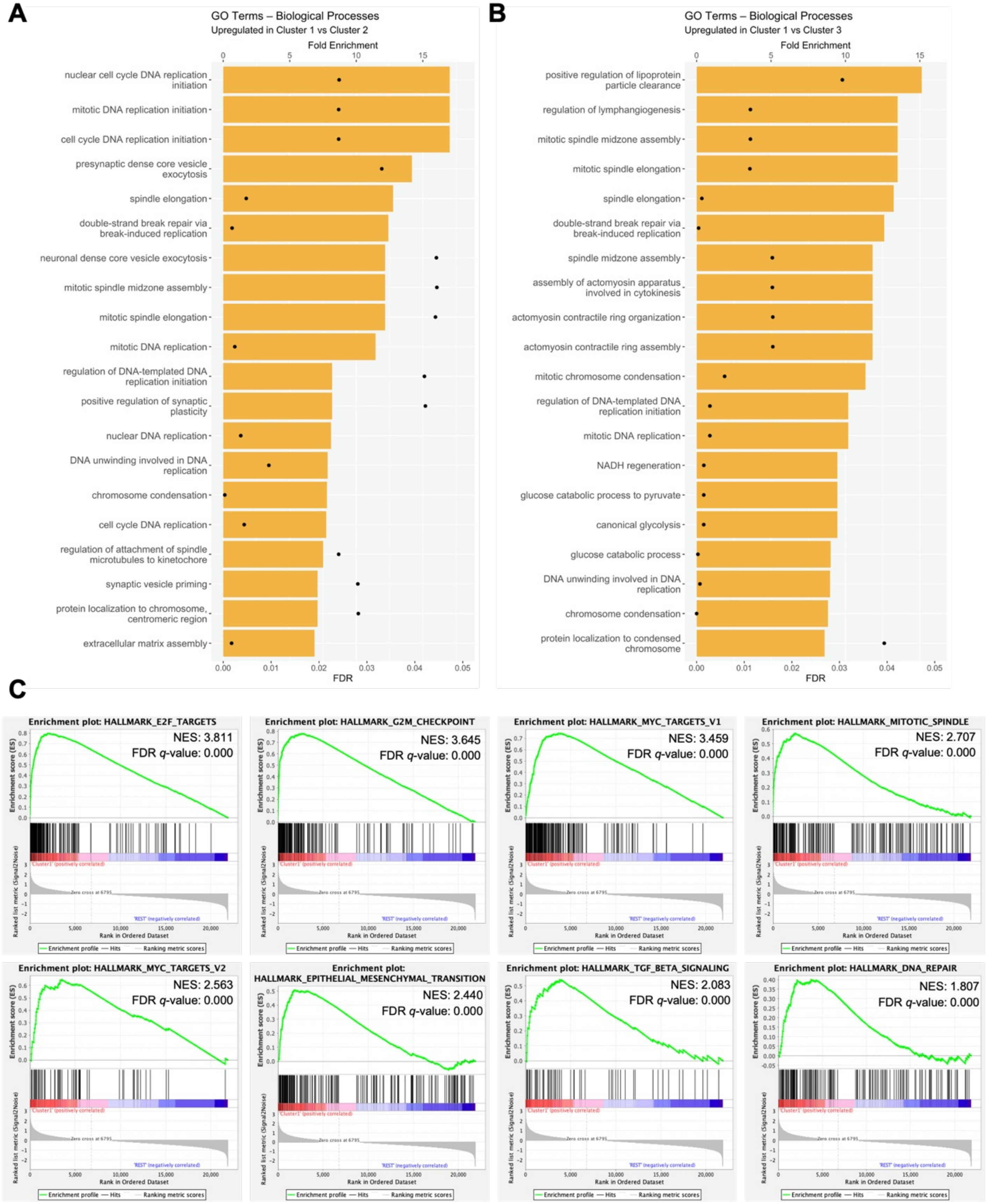

**Supplementary Figure 8.** Cluster 2 tumors are upregulated in oxidative phosphorylation and metabolic biological process GO Terms and GSEA hallmark pathways.

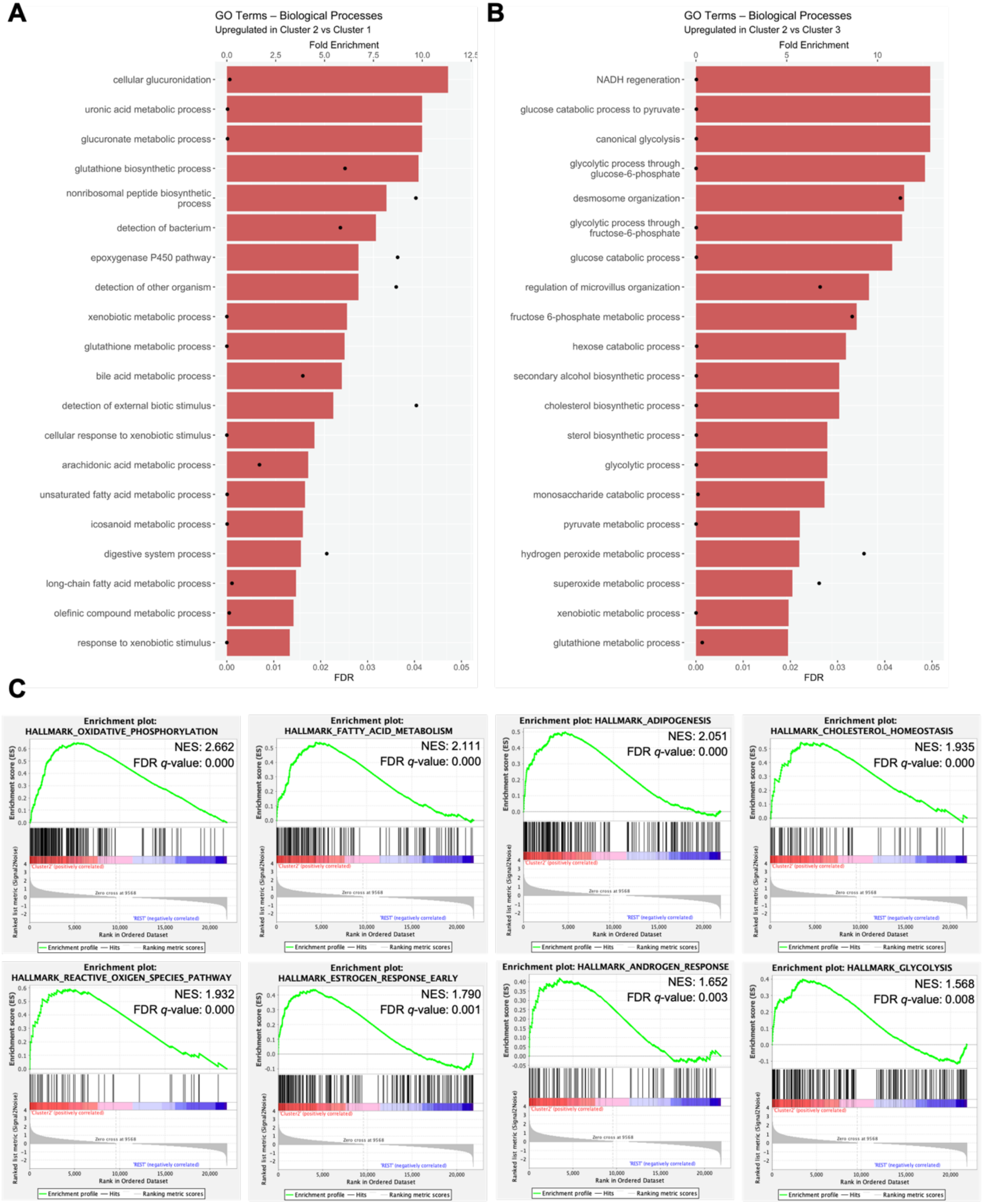

**Supplementary Figure 9.** Cluster 3 tumors are upregulated in immune-modulatory and tyrosine kinase biological process GO Terms and GSEA hallmark pathways.

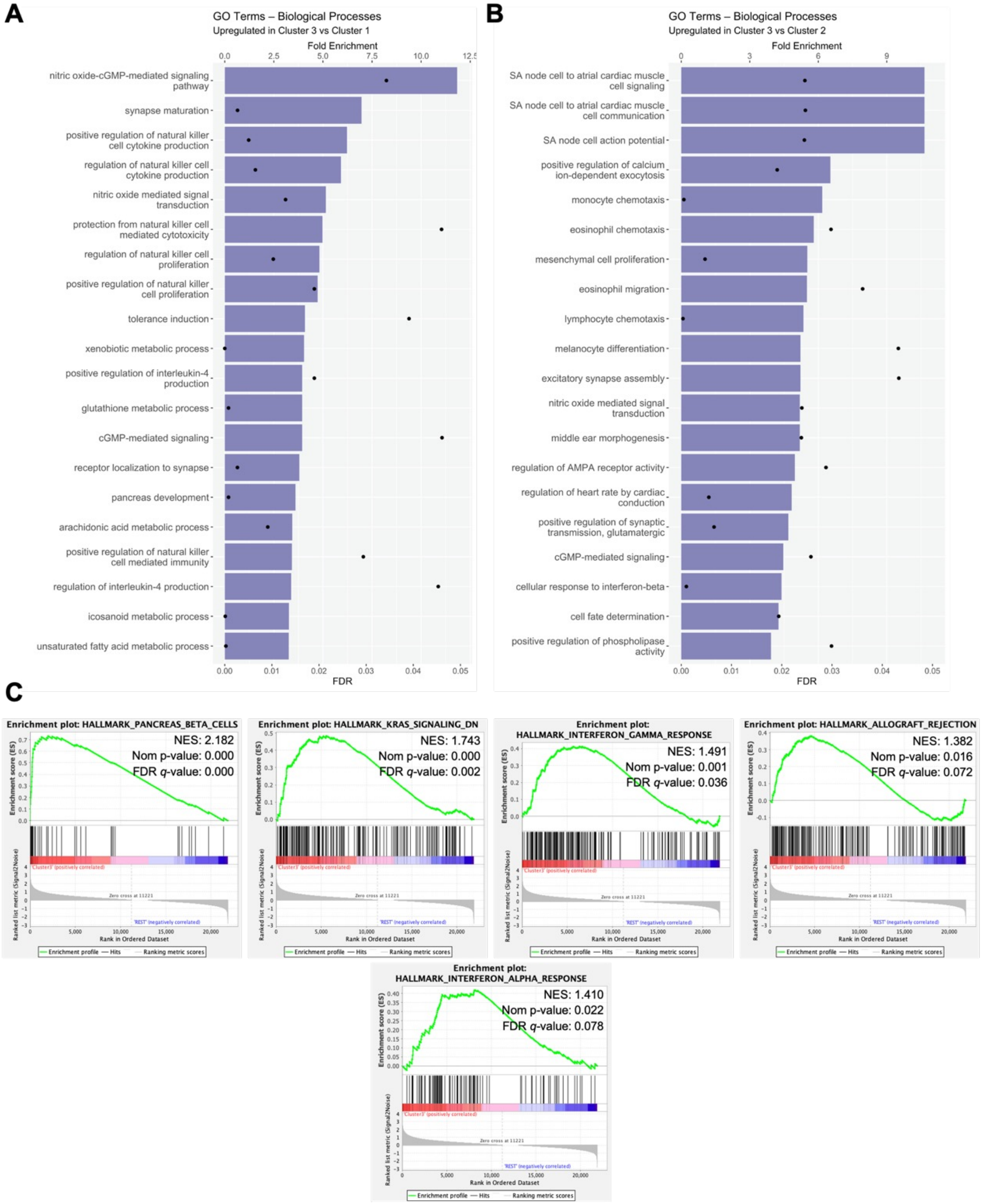

### Supplementary Table 5

| Autochthonous Tumor Endpoint Data | | | Omics Data Availability | | | *In vivo* (orthotopic model) | | | |
|---|---|---|---|---|---|---|---|---|---|
| Mouse / Cell Line ID | Sex | Tumor subtype | RNAseq | WGS | WES | Growth consistency | Liver metastatic frequency | Gemcitabine/ Abraxane sensitivity | FOLFIRINOX sensitivity |
| 011T | M | N/A | - | - | - | Not assessed | Not assessed | Not assessed | Not assessed |
| 106L | F | N/A | - | - | - | Not assessed | Not assessed | Not assessed | Not assessed |
| 171J | M | Squamous | + | + | + | Consistent | 73% | Resistant | Resistant |
| 255J | F | Squamous | + | + | + | Inconsistent | 50% | Not assessed | Not assessed |
| 301J | M | Pancreatic Progenitor | + | + | + | Consistent | 40% | Sensitive | Not assessed |
| 303J | M | Pancreatic Progenitor | + | + | + | Consistent | 40% | Not assessed | Not assessed |
| 314J | M | Squamous | + | + | + | Consistent | 76% | Resistant | Resistant |
| 808J | F | N/A | - | - | - | Not assessed | Not assessed | Not assessed | Not assessed |

| Line | Sex | Subtype | *In vitro* growth | Liver Metastatic Frequency | Gem/Abraxane | FOLFIRINOX |
|---|---|---|---|---|---|---|
| 171J | M | Squamous (Basal-like) | Consistent | 75-100% | Resistant | Resistant |
| 255J | F | Squamous (Basal-like) | Inconsistent (Immune Sensitive) | 50% | Not Assessed | Not Assessed |
| 301J | M | Panc Progenitor (Classical) | Consistent | 40% | Sensitive | Not Assessed |
| 303J | M | Panc Progenitor | Consistent | 40% | Not Assessed | Not Assessed |
| 314J | M | Squamous | Consistent | 76% | Resistant | Resistant |
| 011T | M | Not Assessed | Not Assessed | Not Assessed | Not Assessed | Not Assessed |
| 106L | F | Not Assessed | Not Assessed | Not Assessed | Not Assessed | Not Assessed |
| Z682 | | | | | | |
| Z693 | | | | | | |
| Z696 | | | | | | |

### Supplementary Figure 10:

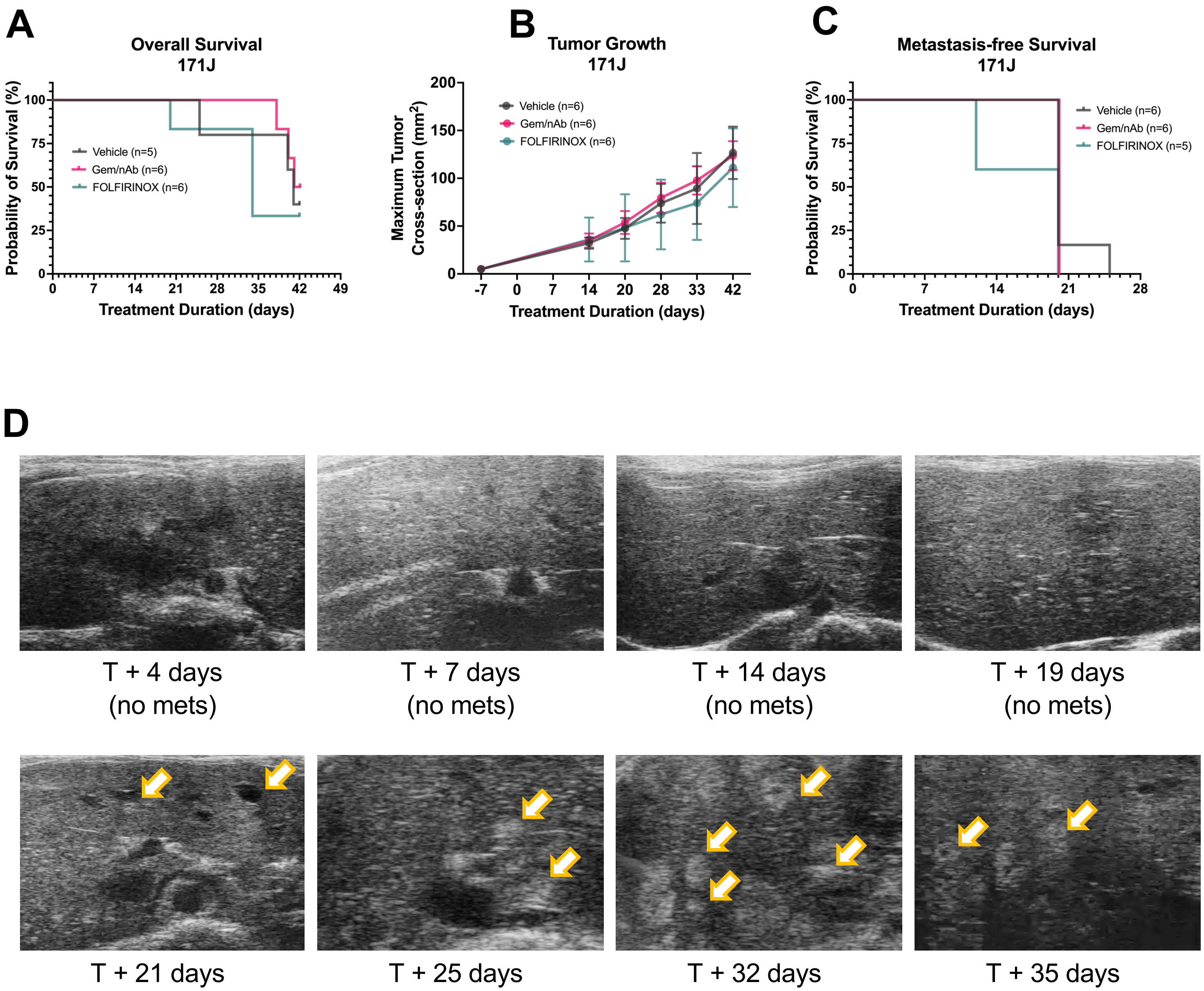
